# Variability in the analysis of a single neuroimaging dataset by many teams

**DOI:** 10.1101/843193

**Authors:** Rotem Botvinik-Nezer, Felix Holzmeister, Colin F. Camerer, Anna Dreber, Juergen Huber, Magnus Johannesson, Michael Kirchler, Roni Iwanir, Jeanette A. Mumford, Alison Adcock, Paolo Avesani, Blazej Baczkowski, Aahana Bajracharya, Leah Bakst, Sheryl Ball, Marco Barilari, Nadège Bault, Derek Beaton, Julia Beitner, Roland Benoit, Ruud Berkers, Jamil Bhanji, Bharat Biswal, Sebastian Bobadilla-Suarez, Tiago Bortolini, Katherine Bottenhorn, Alexander Bowring, Senne Braem, Hayley Brooks, Emily Brudner, Cristian Calderon, Julia Camilleri, Jaime Castrellon, Luca Cecchetti, Edna Cieslik, Zachary Cole, Olivier Collignon, Robert Cox, William Cunningham, Stefan Czoschke, Kamalaker Dadi, Charles Davis, Alberto De Luca, Mauricio Delgado, Lysia Demetriou, Jeffrey Dennison, Xin Di, Erin Dickie, Ekaterina Dobryakova, Claire Donnat, Juergen Dukart, Niall W. Duncan, Joke Durnez, Amr Eed, Simon Eickhoff, Andrew Erhart, Laura Fontanesi, G. Matthew Fricke, Adriana Galvan, Remi Gau, Sarah Genon, Tristan Glatard, Enrico Glerean, Jelle Goeman, Sergej Golowin, Carlos González-García, Krzysztof Gorgolewski, Cheryl Grady, Mikella Green, João Guassi Moreira, Olivia Guest, Shabnam Hakimi, J. Paul Hamilton, Roeland Hancock, Giacomo Handjaras, Bronson Harry, Colin Hawco, Peer Herholz, Gabrielle Herman, Stephan Heunis, Felix Hoffstaedter, Jeremy Hogeveen, Susan Holmes, Chuan-Peng Hu, Scott Huettel, Matthew Hughes, Vittorio Iacovella, Alexandru Iordan, Peder Isager, Ayse Ilkay Isik, Andrew Jahn, Matthew Johnson, Tom Johnstone, Michael Joseph, Anthony Juliano, Joseph Kable, Michalis Kassinopoulos, Cemal Koba, Xiang-Zhen Kong, Timothy Koscik, Nuri Erkut Kucukboyaci, Brice Kuhl, Sebastian Kupek, Angela Laird, Claus Lamm, Robert Langner, Nina Lauharatanahirun, Hongmi Lee, Sangil Lee, Alexander Leemans, Andrea Leo, Elise Lesage, Flora Li, Monica Li, Phui Cheng Lim, Evan Lintz, Schuyler Liphardt, Annabel Losecaat Vermeer, Bradley Love, Michael Mack, Norberto Malpica, Theo Marins, Camille Maumet, Kelsey McDonald, Joseph McGuire, Helena Melero, Adriana Méndez Leal, Benjamin Meyer, Kristin Meyer, Paul Mihai, Georgios Mitsis, Jorge Moll, Dylan Nielson, Gustav Nilsonne, Michael Notter, Emanuele Olivetti, Adrian Onicas, Paolo Papale, Kaustubh Patil, Jonathan E. Peelle, Alexandre Pérez, Doris Pischedda, Jean-Baptiste Poline, Yanina Prystauka, Shruti Ray, Patricia Reuter-Lorenz, Richard Reynolds, Emiliano Ricciardi, Jenny Rieck, Anais Rodriguez-Thompson, Anthony Romyn, Taylor Salo, Gregory Samanez-Larkin, Emilio Sanz-Morales, Margaret Schlichting, Douglas Schultz, Qiang Shen, Margaret Sheridan, Fu Shiguang, Jennifer Silvers, Kenny Skagerlund, Alec Smith, David Smith, Peter Sokol-Hessner, Simon Steinkamp, Sarah Tashjian, Bertrand Thirion, John Thorp, Gustav Tinghög, Loreen Tisdall, Steven Tompson, Claudio Toro-Serey, Juan Torre, Leonardo Tozzi, Vuong Truong, Luca Turella, Anna E. van’t Veer, Tom Verguts, Jean Vettel, Sagana Vijayarajah, Khoi Vo, Matthew Wall, Wouter D. Weeda, Susanne Weis, David White, David Wisniewski, Alba Xifra-Porxas, Emily Yearling, Sangsuk Yoon, Rui Yuan, Kenneth Yuen, Lei Zhang, Xu Zhang, Joshua Zosky, Thomas E. Nichols, Russell A. Poldrack, Tom Schonberg

## Abstract

Data analysis workflows in many scientific domains have become increasingly complex and flexible. To assess the impact of this flexibility on functional magnetic resonance imaging (fMRI) results, the same dataset was independently analyzed by 70 teams, testing nine ex-ante hypotheses. The flexibility of analytic approaches is exemplified by the fact that no two teams chose identical workflows to analyze the data. This flexibility resulted in sizeable variation in hypothesis test results, even for teams whose statistical maps were highly correlated at intermediate stages of their analysis pipeline. Variation in reported results was related to several aspects of analysis methodology. Importantly, meta-analytic approaches that aggregated information across teams yielded significant consensus in activated regions across teams. Furthermore, prediction markets of researchers in the field revealed an overestimation of the likelihood of significant findings, even by researchers with direct knowledge of the dataset. Our findings show that analytic flexibility can have substantial effects on scientific conclusions, and demonstrate factors related to variability in fMRI. The results emphasize the importance of validating and sharing complex analysis workflows, and demonstrate the need for multiple analyses of the same data. Potential approaches to mitigate issues related to analytical variability are discussed.

---

Data analysis workflows in many areas of science have become exceedingly complex, with a large number of processing and analysis steps that involve many possible choices at each of those steps (i.e., “researcher’s degrees of freedom” ^1,2^). There is often no unique correct or “gold standard” workflow, as different options will reflect different tradeoffs and statistical philosophies. Simulation studies have shown that these differences in analytic choices can have substantial effects on results ^3^, but it has not been clear to what degree such variability exists and how it affects reported scientific conclusions in practice. Recent work in psychology has attempted to address this through a “many analysts” approach ^4^, in which the same dataset was analyzed by a large number of groups, uncovering substantial variability in behavioral results across analysis teams. In the Neuroimaging Analysis Replication and Prediction Study (NARPS; www.narps.info), we applied a similar approach to the domain of functional magnetic resonance imaging (fMRI), where analysis workflows are complex and highly variable. Seventy independently acting teams of researchers analyzed the the same functional neuroimaging dataset to test the same nine ex-ante hypotheses. We tested variability in the results across teams and examined the aspects of analysis workflows that were related to this variability. We further employed the novel approach of prediction markets, where participants traded on the outcomes of these analyses, to assess the accuracy of predictions made by researchers in the field ^5–7^.

## Variability of results across analysis teams

The first aim of NARPS was to assess the real-world variability of results across independent analysis teams analyzing the same dataset. The dataset included fMRI data from 108 individuals, each performing one of two versions of a mixed gambles task previously used to study decision-making under risk ^8^. The two versions of the task were designed to address an ongoing debate in the literature regarding the impact of distributions of potential gains/losses on neural activity in this task ^9,10^. A full description of the experimental procedures, validations and the dataset is available in a Data Descriptor ^11^; the dataset is openly available via OpenNeuro at DOI:10.18112/openneuro.ds001734.v1.0.4. Fully reproducible code for all analyses of the data reported here are available at DOI:10.5281/zenodo.3528171.

Neuroimaging researchers were solicited via social media and at the 2018 annual meeting of The Society for Neuroeconomics to participate in the analysis of this dataset. Seventy analysis teams participated in the study. The teams were provided with the raw data, organized according to the Brain Imaging Data Structure (BIDS) ^12^, as well as optional preprocessed data (processed with fMRIprep ^13^). They were asked to analyze the data to test nine ex-ante hypotheses (Table 1), each of which consisted of a description of significant activity in a specific brain region in relation to a particular feature of the experimental design. They were given up to 100 days (varying based on the date they joined) to analyze the data and report for each hypothesis whether it was supported based on a whole-brain corrected analysis (yes / no). In addition, each team submitted a full report of the analysis methods they had used (following COBIDAS guidelines ^14^) and created a collection on NeuroVault ^15^ with one unthresholded and one thresholded statistical map supporting each hypothesis test. To measure variability of results in an ecological manner, the only instructions given to the teams were to perform the analysis as they usually would in their own research groups and to report the binary decision based on their own criteria for a whole-brain corrected result for the specific region described in the hypotheses.

**Table 1.** Hypotheses and results. Each hypothesis is described along with the fraction of teams reporting a whole-brain corrected significant result and two measures reported by the analysis teams for the specific hypothesis (both rated 1-10): (1) How confident are you about this result? (2) How similar do you think your result is to the other analysis teams? For these ordinal measures, median values are presented along with the median absolute deviation in brackets. See Supplementary Materials for analysis of the confidence level and similarity estimation.

|  | Hypothesis description | Fraction of teams reporting a significant result | Median confidence level | Median similarity estimation |
| --- | --- | --- | --- | --- |
| #1 | Positive parametric effect of gains in the vmPFC (equal indifference group) | 0.371 | 7<br>(2) | 7<br>(1.5) |
| #2 | Positive parametric effect of gains in the vmPFC (equal range group) | 0.214 | 7<br>(1.5) | 7<br>(1) |
| #3 | Positive parametric effect of gains in the ventral striatum (equal indifference group) | 0.229 | 6<br>(1) | 7<br>(1) |
| #4 | Positive parametric effect of gains in the ventral striatum (equal range group) | 0.329 | 6<br>(1) | 7<br>(1) |
| #5 | Negative parametric effect of losses in the vmPFC (equal indifference group) | 0.843 | 8<br>(1) | 8<br>(1) |
| #6 | Negative parametric effect of losses in the vmPFC (equal range group) | 0.329 | 7<br>(1) | 7<br>(1) |
| #7 | Positive parametric effect of losses in the amygdala (equal indifference group) | 0.057 | 7<br>(1) | 8<br>(1) |
| #8 | Positive parametric effect of losses in the amygdala (equal range group) | 0.057 | 7<br>(1) | 8<br>(1) |
| #9 | Greater positive response to losses in amygdala for equal range group vs. equal indifference group | 0.057 | 6<br>(1) | 7<br>(1) |

Although teams were not explicitly required to demonstrate expertise in fMRI analysis, members for 69 of the 70 teams had prior publications using fMRI. The dataset, as well as all reports and collections, were kept private until after the prediction markets were closed. The results reported by all teams are presented in Supplementary Table 1. A table describing the methods used by the analysis teams is available with the analysis code. NeuroVault collections containing the submitted statistical maps are available via the links provided in Supplementary Table 2.

The fraction of teams reporting a significant result for each hypothesis is presented in Table 1. Overall, the rates of reported significant findings varied across hypotheses. Only one hypothesis (#5) showed a high rate of significant findings, with 84.3% of teams reporting a significant result. Three other hypotheses showed consistent non-significant findings across groups, with only 5.7% of teams reporting significant findings for hypotheses #7, #8 and #9 (all of which centered on loss-related activity in the amygdala). For the remaining five hypotheses, the results were more variable, ranging from 21.4% to 37.1% of teams reporting a significant result. The extent of the variation in results across teams can be measured as the fraction of teams reporting a different result than the majority of teams (i.e. the absolute distance from consensus). On average across the 9 hypotheses, 20% of teams reported a result that differs from the majority of teams. This is a sizeable variation across teams on average, given that the maximum possible variation is 50%. This implies that the observed fraction of 20% divergent results falls midway between complete consistency in results across teams and completely random results, demonstrating that analytic choices crucially affect reported results.

## Factors related to analytic variability

To better understand the sources of the analytic variability found in the reported binary results, we analyzed the analysis pipelines used by the teams as well as the unthresholded and thresholded statistical maps they provided. There were no two teams with identical analysis pipelines. One team was excluded from all analyses since their reported results were not based on a whole-brain analysis as instructed. Of the remaining 69 teams, thresholded maps of 65 teams and unthresholded (z / t) maps of 64 teams were included in the analyses (see Supplementary Table 3 for detailed reasons for exclusion of the other teams).

### Variability of reported results

We conducted exploratory analyses of the relation between reported hypothesis outcomes and a subset of specific measurable analytic choices and image features. There were several primary sources of analytic variability across teams. First, teams differed in the way they modelled the hypotheses tests (i.e. the regressors and contrasts they included in the model). Second, there were multiple different software packages used. Third, teams differed in the preprocessing steps applied as well as the parameters and techniques used at each preprocessing step. Fourth, teams differed in the threshold used to identify significant effects at each voxel in the brain and the method used to correct for multiple comparisons. Finally, teams differed in how the anatomical regions of interest (ROIs) were defined to determine whether there was a significant effect in each a priori ROI.

A set of mixed effects logistic regression models (with data from N = 64 teams) identified a number of factors that impacted these outcomes (see Supplementary Table 4). The strongest factor was spatial smoothness; higher estimated smoothness of the statistical images (estimated based on the unthresholded statistical maps using FSL’s smoothest function) was associated with a greater likelihood of significant outcomes (*p* < 0.001, delta pseudo-*R*^*2*^ = 0.04; mean FWHM 9.69 mm, range 2.50 - 21.28 mm across teams). Interestingly, while estimated smoothness was related to the width of the applied smoothing kernel (*r* = 0.71; median applied smoothing 5 mm, range 0 - 9 mm across teams), the applied smoothing value itself was not significantly related to positive outcomes in a separate analysis, suggesting that the relevant smoothness may have arisen from other analytic steps in addition to explicit smoothing on its own. In particular, exploratory analyses showed that the inclusion of head motion parameters in the statistical model was associated with lower image smoothness (*p* = 0.014). An effect on decision outcomes was also found for the software package used (*p* = 0.004, delta pseudo-*R*^*2*^ = 0.04; N = 23 [SPM], 21 [FSL], 7 [AFNI], 13 [Other]), with the FSL package being associated with a higher likelihood of significant results across all hypotheses compared to the SPM package; odds ratio = 6.69), and for the effect of different multiple test correction methods (*p* = 0.024, delta pseudo-*R*^*2*^ = 0.02: N = 48 [parametric], 14 [nonparametric], 2 [other]), with parametric correction methods leading to higher rates of detection than nonparametric methods. No significant effect on the decision outcomes was detected for the use of standardized preprocessed data (with fMRIprep) versus custom preprocessing pipelines (48% of included teams used fMRIprep; *p* = 0.132) or the inclusion of head motion parameters in the statistical model (used by 73% of the teams; *p* = 0.281).

### Variability of thresholded statistical maps

The nature of analytic variability across the whole brain was further explored by analyzing the statistical maps submitted by the teams. The thresholded activation maps were highly sparse (median number of activated voxels over teams ranged from 167 to 9,383 across hypotheses, out of 228,483 voxels in the MNI standard mask). Binary agreement between thresholded maps over all voxels was relatively high (median percent agreement ranged from 93% to 99% across hypotheses), largely reflecting agreement on which voxels were never active. However, when restricted only to voxels showing any activation over teams, overlap was very low (median similarity ranging from 0.00 to 0.06 across hypotheses). This may have reflected the substantial variability in the number of activated voxels found by each team; for every hypothesis, the number of voxels found as active ranged across teams from zero to tens of thousands (Supplementary Table 5). Analysis of overlap between activated voxels was consistent with the variability in the reported hypothesis results, with most voxels in the thresholded maps showing inconsistent binary values. The maximum proportion of teams with activation in any single voxel for a given hypothesis was 0.77 (range 0.23 - 0.77; Figure 1). However, a coordinate-based meta-analysis using activation likelihood estimation (ALE) ^16,17^ across teams, which imposes additional smoothing, demonstrated convergent patterns of activation for all hypotheses (Supplementary Figure 1). Altogether, analysis of the similarity between thresholded statistical images suggests that these maps are substantially diverse, but aggregating across analyses can yield more consistent results.

**Figure 1.**
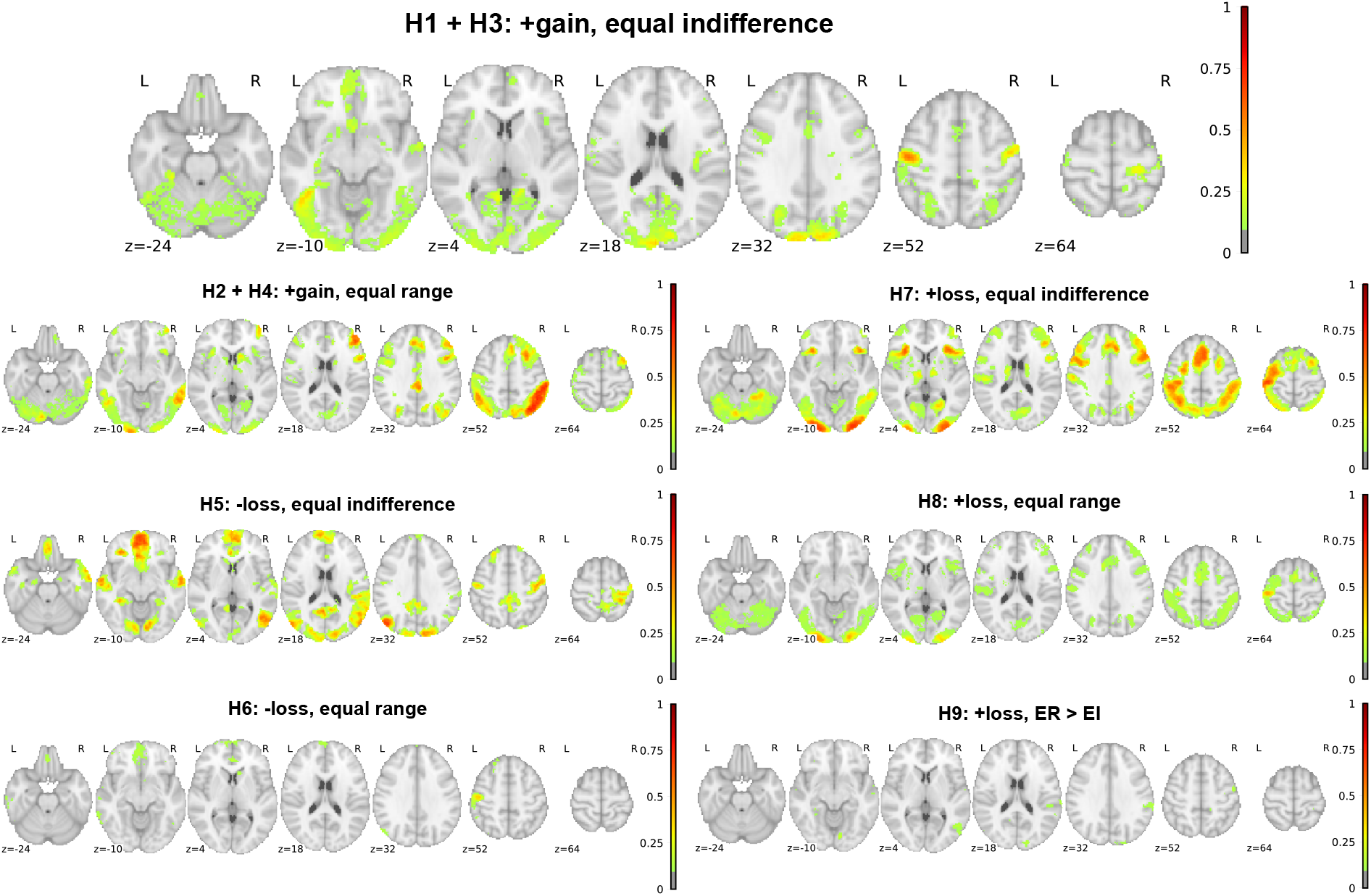
Voxels overlap. Maps showing at each voxel the proportion of teams reporting significant activations in their thresholded statistical map, for each hypothesis (labeled H1 - H9), thresholded at 10% (i.e., voxels with no color were significant in fewer than 10% of teams). +/− refers to direction of effect, gain/loss refers to the effect being tested, and equal indifference (EI) / equal range (ER) refers to the group being examined or compared. Hypotheses #1 and #3, as well as hypotheses #2 and #4, share the same statistical maps as the hypotheses are for the same contrast and experimental group, but for different regions (see Table 1). Images can be viewed at https://identifiers.org/neurovault.collection:6047

### Variability of unthresholded statistical maps

Analysis of correlation between unthresholded Z-statistic maps across teams demonstrated that for each hypothesis, there was a large cluster of teams whose statistical maps were strongly positively correlated with one another (see Figure 2 for an example with hypothesis 1, and Supplementary Figures 2-7 for other hypotheses). Overall correlation (mean Spearman correlation) between pairs of unthresholded maps was moderate (mean correlation range 0.18 - 0.52 across hypotheses), with higher correlations within the main cluster of analysis teams (range 0.44 - 0.85 across hypotheses) (see Supplementary Table 6). Correlations between the unthresholded maps were further assessed by modeling the median Spearman correlation of each team with the average pattern across teams as a function of analysis method using linear regression. Estimated spatial smoothness of the statistical images (averaged across hypotheses) was significantly associated with correlation with the mean pattern (*p* = 0.023, delta *r*^*2*^ = 0.07), as was the use of movement modeling (*p* = 0.021, delta *r*^*2*^ = 0.08).

**Figure 2.**
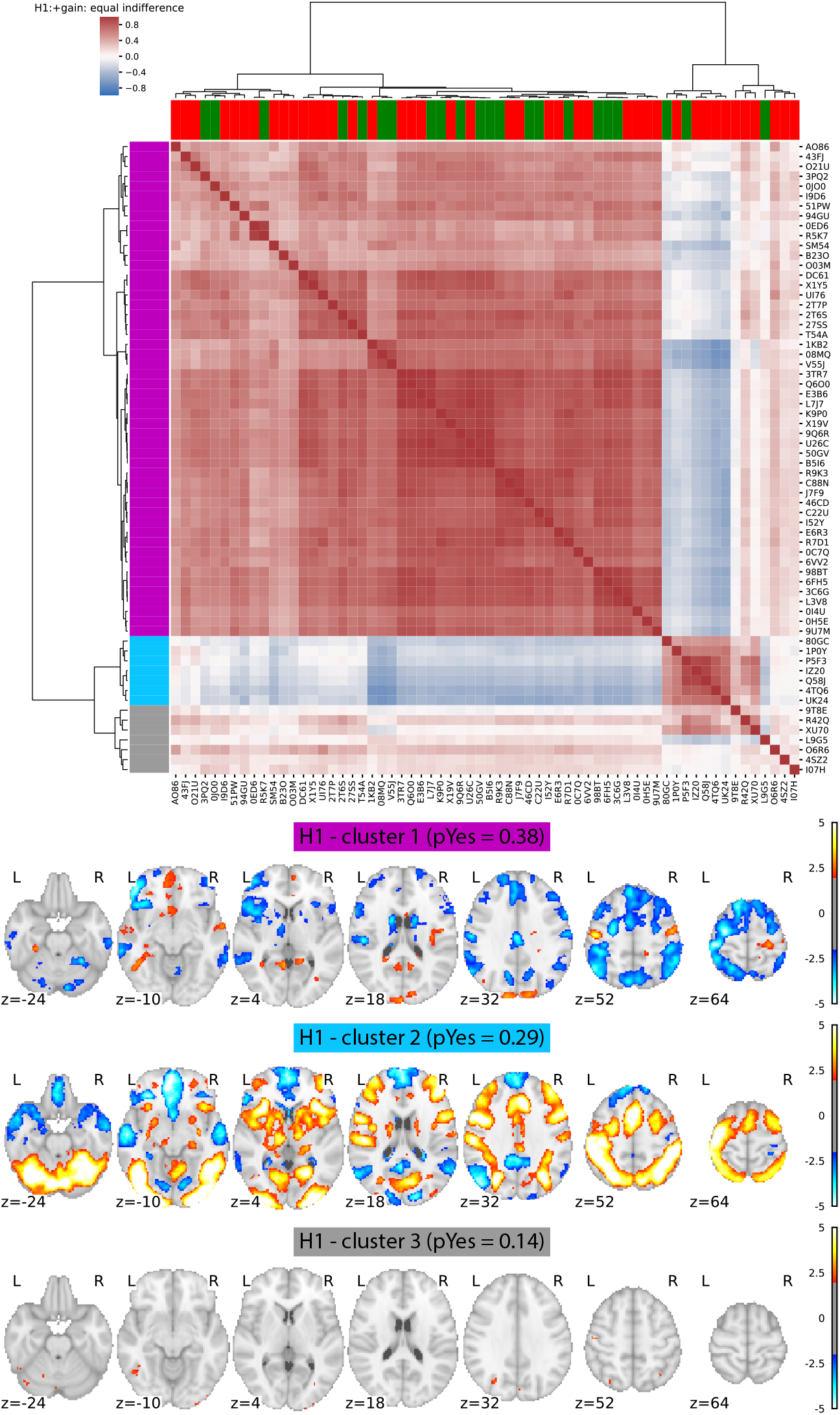
Analytic variability in whole-brain statistical results for Hypothesis 1. **Top panel**: Spearman correlation values between whole-brain unthresholded statistical values for each team were computed and clustered according to their similarity (using Ward clustering on Euclidean distances). Row colors (left) denote cluster membership, while column colors (top) represent hypothesis decisions (green: Yes, red: No). Brackets represent clustering. **Bottom panel**: Average statistical maps (thresholded at uncorrected z > 2.0) for each of the three clusters depicted in panel A. The probability of reporting a positive hypothesis outcome is presented for each cluster. Images can be viewed at https://identifiers.org/neurovault.collection:6048.

Variability across unthresholded statistical maps was assessed by computing the between- teams variability using an equivalent of the tau-squared statistic commonly used to assess heterogeneity in meta-analysis; in the case where all results are equivalent, this statistic should take a value approaching zero. Median tau across teams was well above one (range across hypotheses: 1.13-1.85), and visualization of voxelwise tau maps (Supplementary Figure 8) showed much higher variability in activated regions, with some voxels showing values greater than 5. As a point of comparison, the sampling variability of T-scores over *different datasets* always has a standard deviation of at least 1.0, and thus it is notable that inter-team variability on the same dataset is often substantially larger.

For Hypotheses #1 and #3, there was also a subset of seven teams whose unthresholded maps were anticorrelated with those of the main cluster of teams. A comparison of the average map for the anticorrelated cluster for Hypotheses #1 and #3 confirmed that this map was highly correlated (*r* = 0.87) with the overall task activation map (averaged across the relevant group for these hypotheses) as reported in the NARPS Data Descriptor^11^. Further analysis of the model specifications for the six teams with available modeling details showed that four of them appeared to use models that did not properly separate the parametric effect of gain from overall task activation; because of the general anticorrelation of value system activations with task activations^18^, this model mis-specification led to an anticorrelation with the parametric effects of gain. In the other two cases, the model included multiple regressors that were correlated with the gain parameter, which drastically modified the interpretation of the primary gains regressor.

The apparent discrepancy between overall- vxc correlations of unthresholded maps and divergence of reported binary results (even within the highly correlated main cluster) suggested that the variability in regional hypothesis test results might be due to procedures related to statistical correction for multiple comparisons and anatomical specification of the region of interest. To further assess this, we applied a consistent thresholding and correction method and anatomical region specification on the unthresholded maps across all teams for each hypothesis. This analysis showed that even using a correction method known to be liberal and a standard anatomical definition for all regions, the degree of variability across results was qualitatively similar to that of the reported hypothesis decisions (Supplementary Table 7 and Supplementary Figure 9).

Meta-analytic approaches that aggregate information across analyses are one potential solution to the issue of analytic variability. We assessed the consistency across teams using an image-based meta-analysis (accounting for correlations due to common data), which demonstrated significant active voxels for all hypotheses except for #9 after false discovery rate correction (see Supplementary Figure 10) and confirmatory evidence for Hypotheses 2, 4, 5, and 6. These results confirmed the coordinate-based meta-analysis reported above (Supplementary Figure 1) in showing that relatively inconsistent results at the individual team level underlie consistent results when the team’s results are combined.

## Prediction markets

The second aim of NARPS was to test whether peers in the field could predict the results obtained in aggregate by the analysis teams using prediction markets. Prediction markets are assumed to aggregate private information distributed among traders, and can generate and disseminate a consensus among market participants. Hanson^19^ first suggested that prediction markets could be a potentially important tool for assessing scientific hypotheses. Recent studies that used prediction markets to predict the replicability of experimental results in the social sciences have yielded promising results^5–7,20^. Predictions revealed by market prices were correlated with actual replication outcomes, although with a tendency towards overestimating the replicability of findings in several studies^5–7,20^.

In NARPS, we ran two separate prediction markets: one involving members from the analysis teams (“team members” prediction market) and an additional independent market for researchers in the field who had not participated in the analysis (“non-team members” prediction market). The prediction markets were open for 10 consecutive days approximately 1.5 months after all analysis teams had submitted their results (which were kept private and confidential). On each market, traders were endowed with tokens worth $50 and traded via an online market platform on the main outcome measures of the fMRI analyses, i.e., the fraction of teams reporting a significant result for each hypothesis. The prediction market prices serve as measures of the aggregate beliefs of traders for the fraction of teams reporting a significant result for each hypothesis. Overall, *n* = 65 traders actively traded in the “non-team members” prediction market and *n* = 83 traders actively traded in the “team members” prediction market. After the prediction markets closed, traders were paid based on their performance in the markets. The analysis of the prediction markets was pre-registered on OSF (https://osf.io/59ksz/). Note that since most of the analyses were performed on the final market prices (i.e., the markets’ predictions), for which there is one value per hypothesis per market, the number of observations for each set of prediction markets was low (*N* = 9), leading to very limited statistical power. Therefore, the results should be interpreted cautiously.

The predictions (i.e., the final market prices) ranged from 0.073 to 0.952 (*m* = 0.599, *sd* = 0.325) in the “team members” prediction market and from 0.476 to 0.882 (*m* = 0.690, *sd* = 0.137) in the “non-team members” prediction market. Except for the prediction of a single hypothesis (Hypothesis #7) in the “team members” set of markets, all predictions were outside the 95% confidence intervals of the fundamental values (i.e. the proportion of teams reporting a significant result for each hypothesis; see Figure 3 and Supplementary Table 8 for details).

**Figure 3:**
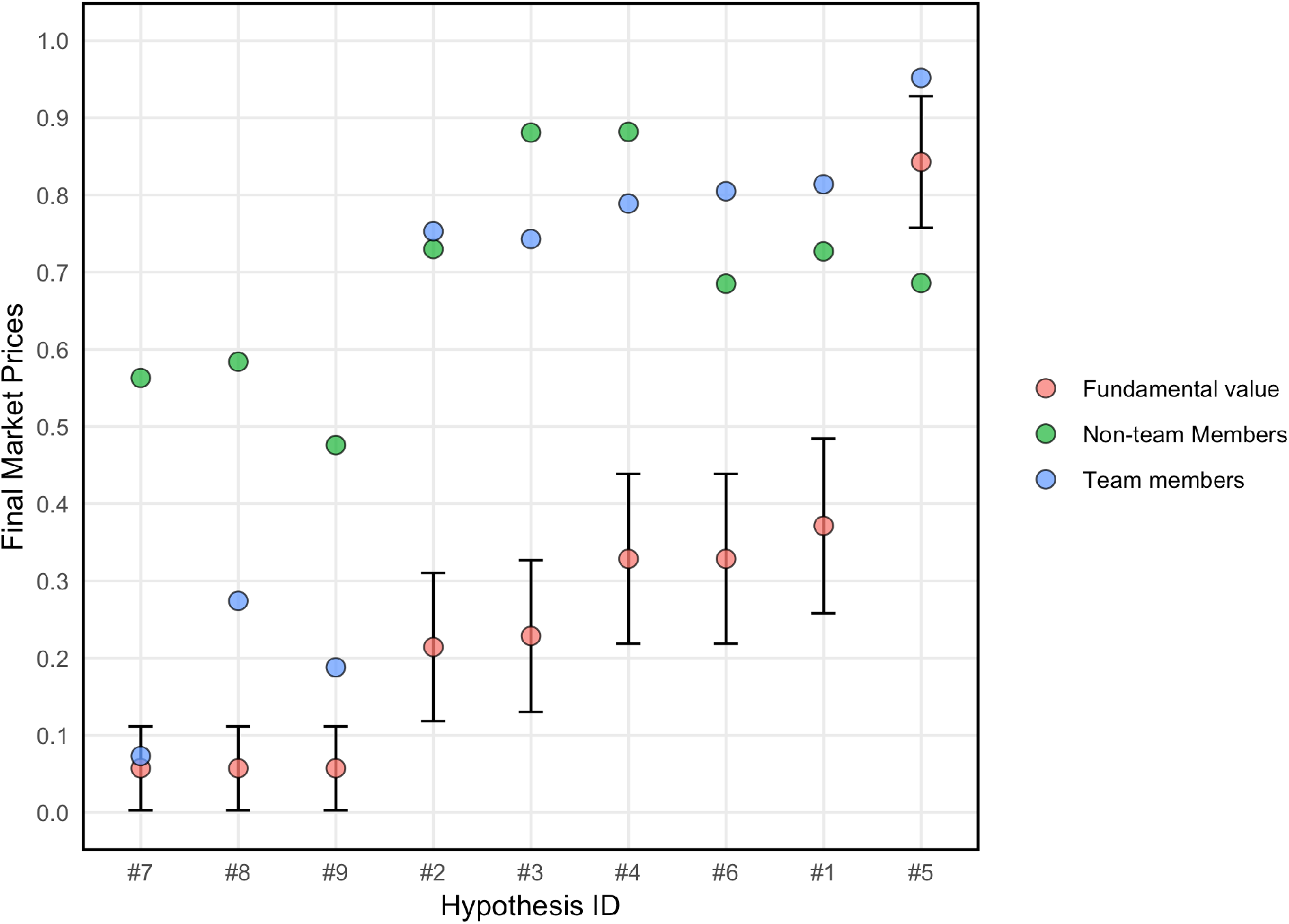
Prediction market beliefs. The figure depicts final market prices (i.e., aggregated market beliefs) for the “team members” (blue dots) and the “non-team members” (green dots) prediction markets as well as the observed fraction of teams reporting significant results, i.e., the fundamental value (pink dots), and the corresponding 95% confidence intervals for each of the nine hypotheses. Confidence intervals were constructed by assuming convergence of the binomial distribution towards the normal.

For the “team members” prediction market, the Spearman (rank-order) correlation between final market’s predictions and fundamental values was 0.962 (*p* < 0.001, *n* = 9). The Spearman correlation between the market’s predictions in the “non-team members” set of markets and the fundamental values was 0.553 (*p* = 0.122, *n* = 9). The Spearman correlation coefficient between the market’s predictions in the “team members” and “non-team members” set of markets was 0.500, but it was not statistically significant (*p* = 0.170, *n* = 9). Wilcoxon signed-rank tests suggest that both “team members” and “non-team members” systematically overestimated the actual fraction of analysis teams reporting significant results for the ex-ante hypotheses (“team members”: *z* = 2.886, *p* = 0.004, *n* = 9; “non-team members”: *z* = 2.660, *p* = 0.008, *n* = 9). The mean absolute error was 0.323 (*sd* = 0.203, *min* = 0.016, *max* = 0.539) for “team members” and 0.449 (*sd* = 0.146, *min* = 0.157, *max* = 0.652) for “non-team members”. The result in the “team members” prediction market was not driven by over-representation of teams reporting significant results (see Supplementary Materials: Supplementary Figure 11 and Prediction markets results/exploratory analyses). Market’s predictions in the “team members” prediction market did not significantly differ from those of the “non-team members” prediction markets (Wilcoxon signed-rank test, *z* = 1.035, *p* = 0.301, *n* = 9), but as mentioned above, the statistical power for this test was limited. Team members generally traded in the direction consistent with their own team’s results (Supplementary Table 9), which may explain why the collective market’s predictions were more accurate than those of non-team members (see Figure 3). For additional results of the prediction markets see Supplementary Materials.

## Discussion

The analysis of a single functional neuroimaging dataset by 70 independent analysis teams revealed substantial variability in reported binary results, with high levels of disagreement across teams of their outcomes on a majority of tested pre-defined hypotheses. Each team used a different analysis pipeline. For Hypotheses 7-9 with the lowest rate of endorsement one could still find four different analysis pipelines used in practice by research groups in the field that resulted in a significant outcome, and for other hypotheses that number was even greater. As nearly every paper in many scientific disciplines (including neuroimaging) currently publishes results based on null hypothesis significance testing (NHST), our findings highlight the fact that it is hard to estimate the reproducibility of single studies that are performed using one single analysis pipeline. Importantly, analyses of the underlying statistical parametric maps on which the inferences were based revealed greater consistency than expected from the reported inferences, with significant consensus in activated regions across groups via two different meta-analytic approaches. As shown in Figure 2, teams with highly correlated underlying statistical maps nonetheless reported highly divergent hypothesis outcomes. Detailed analysis of the workflow descriptions and statistical results submitted by the analysis teams identified several common analytic variables that were related to differential reporting of significant outcomes, including the spatial smoothness of the data (which is the result of multiple factors in addition to the applied smoothing kernel), and choices of analysis software and correction method. In addition, we identified model specification errors for several analysis teams leading to statistical maps that were anticorrelated with the majority of teams. Prediction markets demonstrated that researchers generally overestimated the likelihood of significant results across hypotheses, even those who had analyzed the data themselves, reflecting substantial optimism bias by researchers in the field.

Given the substantial amount of analytic variability we found to be present in practice, leading to substantial variability of reported hypothesis results with the same data, we believe that steps need to be taken to improve the reproducibility of data analysis outcomes. First, as a practical immediate step in the field of neuroimaging we suggest that unthresholded statistical maps should be shared as a standard practice alongside thresholded statistical maps using tools such as NeuroVault ^15^. In the long run, the shared maps will allow the use of image-based meta-analysis, which we found to provide robust results across laboratories. Second, the use of pre-registration ^21^ or registered reports ^22^ can minimize researchers’ degrees of freedom and their effect on neuroimaging results. Finally, publicly sharing data and analysis code should become common practice, to enable others to run their own analysis with the same data or validate the code used. These good practices would not prevent analytic variability as demonstrated here, but would ensure that analysis variables were not selected in a data-dependent manner. All of the data and code for this study are publicly available with a fully reproducible execution environment for all figures and results. We believe that this can serve as an example for future studies.

Foremost, we propose that complex datasets should be analyzed using multiple analysis pipelines, preferably by more than one researcher, who would be blinded to the hypotheses of interest^23^ and to the results obtained using other pipelines. Achieving such “multiverse analysis”^24^ at scale will require the development of fully automated configurable statistical analysis tools (e.g.^25^) that can run a broad range of reasonable pipelines and assess their convergence. Different versions of such “multiverse” analysis have been suggested in other fields^26,27^, but are not widely used. Analysis pipelines should also be validated using simulated data in order to assess their validity with regard to ground truth (as done in the present study for the code used to analyze the variability across teams), and assessed for their effects on predictions with new data ^28^. The present investigation was limited to the analysis of a single functional neuroimaging dataset, but it seems highly likely that similar variability will be present for fields of research where the data are high-dimensional and the analysis workflows are complex and varied. Our findings add new urgency to similar ecologically valid assessments of analytic variability in those fields as well (for additional discussion see Supplementary Discussion).

## Methods

### fMRI dataset

In order to test the variability of neuroimaging results across analysis pipelines used in practice in research laboratories, we distributed a single fMRI dataset to independent analysis groups from around the world, requesting them to test nine pre-defined hypotheses. The full dataset is publicly available on OpenNeuro (DOI: 10.18112/openneuro.ds001734.v1.0.4) and is described in details in a Data Descriptor^11^.

Shortly, the fMRI dataset consisted of data from 108 participants performing a mixed gamble task, a task often used to study decision-making under risk. In this task, participants are asked on each trial to accept or reject a presented prospect. The prospects consist of an equal 50% chance of either gaining a given amount of money or losing another, similar or different, amount of money. Participants were divided into two groups: in the “equal indifference” group (*N* = 54), the potential losses were half the size of the potential gains^8^(reflecting the “loss aversion” phenomenon, where people tend to be more sensitive to losses compared to equal-sized gains^29^); in the “equal range” group (*N* = 54), the potential losses and the potential gains were taken from the same scale^9,10^. The two groups were used to resolve inconsistencies of previous results.

The dataset was distributed to the teams via Globus (https://www.globus.org/). The distributed dataset included raw data of 108 participants (*N* = 54 for each experimental group), as well as the same data after preprocessing with fMRIprep version 1.1.4 [RRID:SCR_016216]^13^. The fMRIprep preprocessing mainly included brain extraction, spatial normalization, surface reconstruction, head motion estimation and susceptibility distortion correction. Both the raw and the preprocessed datasets underwent quality assurance (described in detail in the Data Descriptor^11^).

### Pre-defined hypotheses

Previous studies with the mixed gamble task suggested that activity in the vmPFC and ventral striatum, among other brain regions, is related to the magnitude of the potential gain^8^. A fundamental open question in the field of decision-making under risk is whether the same brain regions also code the magnitude of the potential loss (through negative activation), or rather potential losses are coded by regions related to negative emotions, such as the amygdala ^8–10^. The specific hypotheses included in NARPS were chosen to address this open question, using two different designs that were used in those previous studies (i.e., equal indifference versus equal range). Each analysis team tested nine pre-defined hypotheses (see Table 1). Each hypothesis predicted fMRI activation in a specific brain region, in relation to a specific aspect of the task (gain / loss amount) and a specific group (equal indifference / equal range, or a comparison between the two groups). Therefore, for each hypothesis, the maximal sample size was 54 participants (Hypotheses #1-8) or 54 participants per group in the group comparison (Hypothesis #9). Although the hypotheses referred to specific brain regions, analysis teams were instructed to report their results based on a whole-brain analysis (and not on a region of interest based analysis, as sometimes used in fMRI studies).

### Analysis teams recruitment and instructions

We recruited analysis teams via social media, mainly Twitter and Facebook, as well as during the 2018 annual meeting of The Society for Neuroeconomics. Ninety-seven teams registered to participate in the study. Each team consisted of up to three members. To ensure independent analyses across teams, and to prevent influencing the subsequent prediction markets, all team members signed an electronic nondisclosure agreement that they would not release, publicize, or discuss their results with anyone until the end of the study. All team members of 82 teams signed the nondisclosure form. They were offered co-authorship on the present publication in return for their participation.

Analysis teams were provided with access to the full dataset. They were asked to freely analyze the data with their usual analysis pipeline to test the nine hypotheses and report a binary decision for each hypothesis on whether it was significantly supported based on a whole-brain analysis. While the hypotheses were region specific, we clearly requested a whole-brain analysis result to avoid the need of teams to create and share masks of regions. Each team also filled in a full report of the analysis methods used (following COBIDAS guidelines ^14^) and created a collection on NeuroVault ^15^ [RRID:SCR_003806] with one unthresholded and one thresholded statistical maps for each hypothesis, on which their decisions were based (teams could optionally include additional maps in their collection). For each result (i.e., the binary decision on whether a given hypothesis was supported by the data or not), teams further reported how confident they were in this result and how similar they thought their result was to the results of the other teams (each measure was an integer between 1 [not at all] to 10 [extremely]). These measures are presented in Table 1 and Supplementary Table 1. In order to measure variability of results in an ecological manner, instructions to the analysis teams were minimized and the teams were asked to perform the analysis as they usually do in their own laboratory and to report the binary decision based on their own criteria.

Seventy of the 82 teams submitted their results and reports by the final deadline (March 15th, 2019). The dataset, as well as all reports and collections, were kept private until the end of the study and closure of the prediction markets. In order to avoid identification of the teams, each team was provided with a unique random 4-character team ID.

Overall, 180 participants were part of NARPS analysis teams. Participating teams were located in 17 countries/regions around the world: Australia (3 participants), Austria (3), Belgium (9), Brazil (3), Canada (16), China (2), Finland (1), France (4), Germany (21), Italy (13), the Netherlands (8), Spain (4), Sweden (4), Switzerland (3), Taiwan (3), UK (7) and the USA (76). Out of 70 analysis teams, five teams consisted of one member, 20 teams consisted of two members and 45 teams consisted of three members. Out of the 180 team members, there were 62 principal investigators (PIs), 43 post-doctoral fellows, 53 graduate students and 22 members from other positions (e.g. data scientists or research analysts).

### Data and code availability

Code for all analyses of the reports and statistical maps submitted by the teams is openly shared in GitHub (https://github.com/poldrack/narps). Image analysis code was implemented within a Docker container, with software versions pinned for reproducible execution (https://cloud.docker.com/repository/docker/poldrack/narps-analysis). Python code was automatically tested for quality using the flake8 static analysis tool and the codacy.com code quality assessment tool, and the results of the image analysis workflow were validated using simulated data. Imaging analysis code was independently reviewed by an expert who was not involved in writing the original code. Prediction market analyses were performed using R v3.6.1; packages were installed using the checkpoint package, which reproducibly installs all package versions as of a specified date (8/13/2019). Analyses reported in this manuscript were performed using code release v1.0.1 (DOI: 10.5281/zenodo.3528171).

Reviewers may obtain anonymous access to the data and run the full image analysis stream by following the directions at: https://github.com/poldrack/narps/tree/master/ImageAnalyses.

Access to the raw data requires specifying a URL for the dataset, which is: https://zenodo.org/record/3528329/files/narps_origdata_1.0.tgz

Results (automatically generated figures, results, and output logs) for imaging analyses are available for anonymous download at DOI:10.5281/zenodo.3528320.

Although not required to, several analysis teams also publicly shared their analysis code. Supplementary Table 10 includes these teams along with the link to their code.

### Factors related to analytic variability

In order to explore the factors related to the variability in results across teams, the reports of all teams were manually annotated to create a table describing the methods used by each team.

We performed exploratory analysis of the relation between the reported hypothesis outcomes and several analytic choices and image features using mixed effects logistic regression models implemented in R, with the lme4 package ^30^. The factors included in the model were: Hypothesis number, estimated smoothness (based on FSL’s smoothest function), use of standardized preprocessing, software package, method of correction for multiple comparisons and modeling of head movement. The teams were modeled as a random effect. One team submitted results that were not based on a whole brain analysis as requested, and therefore their data were excluded from all analyses.

In addition, we performed exploratory analyses to explore the variability across statistical maps submitted by the teams. The unthresholded and thresholded statistical maps of all teams were resampled to common space (FSL MNI space, 91×109×91, 2mm isotropic) using nilearn^31^[RRID:SCR_001362]. For unthresholded maps, we used 3rd order spline interpolation; for thresholded maps, we used linear interpolation and then thresholded at 0.5, to prevent artifacts that appeared when using nearest neighbor interpolation. Of the 69 teams included in the analyses, unthresholded maps of five teams and thresholded maps of four teams were excluded from the image-based analyses (see Supplementary Table 3 for details). Since some of the hypotheses reflected negative activations, which can be represented by either positive or negative values in the statistical maps, depending on the model used, we asked the teams to report the direction of the values in their maps for the relevant hypotheses (#5, #6, and #9). Unthresholded maps were corrected to address sign flips for reversed contrasts as reported by the analysis teams. In addition, t values were converted to z values with Hughett's transform^32^. All subsequent analyses of the unthresholded maps were performed only on voxels that contained non-zero data for all teams (range across hypotheses: 111062 - 145521 voxels).

We assessed the agreement between thresholded statistical maps using percent agreement, i.e. the percent of voxels that have the same binary value. Because the thresholded maps are very sparse, these values are necessarily high when computed across all voxels. Therefore, we also computed the agreement between pairs of statistical maps only for voxels that were nonzero for at least one member of each pair. To further test the agreement across teams, we performed a coordinate-based meta-analysis with activation likelihood estimation (ALE) ^16,17^. This analysis was performed with the NIMARE software package [RRID:SCR_017398] using peak locations identified from thresholded maps for each team. Correction for multiple tests was applied using false discovery rate at the 5% threshold^33^.

We further computed the correlation between the unthresholded images of the 64 teams. The correlation matrices were clustered using Ward clustering; the number of clusters was set to three for all hypotheses based on visual examination of the dendrograms. A separate mean statistical map was then created for the teams in each cluster (see Figure 2 and Supplementary Figures 2-7). Drivers of map similarity were further assessed by modeling the median correlation distance of each team from the average pattern as a function of several analysis decisions (e.g. smoothing, whether or not the data preprocessed with fMRIprep were used, etc.).

To assess the impact of variability in thresholding methods and anatomical definitions across teams, unthresholded Z maps for each team were thresholded using a common approach. Z maps for each team were translated to p-values, which were then thresholded using two approaches: a heuristic correction (known to be liberal^34^), and a voxelwise false discovery rate correction. Note that it was not possible to compute the commonly-used familywise error correction using Gaussian random field theory because residual smoothness was not available for each team. We then identified whether there were any suprathreshold voxels within the appropriate anatomical region of interest for each hypothesis. The regions of interest for the ventral striatum and amygdala were defined anatomically based on the Harvard-Oxford anatomical atlas. Since there is no anatomical definition for the ventromedial prefrontal cortex, we defined the region using a conjunction of anatomical regions (including all anatomical regions in the Harvard-Oxford atlas that overlap with the ventromedial portion of the prefrontal cortex) and a meta-analytic map obtained from neurosynth.org^35^ for the search term “ventromedial prefrontal”.

An image-based meta-analysis was used to quantify the evidence for each hypothesis across analysis teams (see Supplementary Figure 10), accounting for the lack of independence due to the use of a common dataset across teams. While there are different meta-analysis-inspired approaches that could be taken (e.g. a random effects meta-analysis that penalizes for inter-team variation), we sought an approach that would preserve the typical characteristics of the teams’ maps. In particular, the meta-analytic statistical map is based on the mean of teams’ statistical maps, but is shifted and scaled by global factors so that the mean and variance are equal to the original image-wise means and variances averaged over teams. Under a complete null hypothesis of no signal anywhere for every team and every voxel, the resulting map can be expected to produce nominal standard normal *z*-scores, and in the presence of signal will reflect a consensus of the different results.

The coordinate-based meta-analysis method is as follows. Let *N* be the number of teams, *μ* be the (scalar) mean over space of each team’s map, averaged over teams, *σ*^2^ likewise the spatial variance averaged over teams, and let **Q** be the *N*×*N* correlation matrix, computed using all voxels in the statistical map. Then let *Z*_*ik*_ be the *z*-value for voxel *i* and team *k*, and *M*_*i*_ the mean of those *N z*-values at voxel *i*. The variance of *M*_*i*_ is *σ*^2^**1**^T^**Q1**/*N*^2^, where **1** is a *N*-vector of ones. We center and standardize *M*_*i*_, and then rescale and shift to produce a meta-analytic *Z-*map with mean *μ* and variance *σ*^2^:

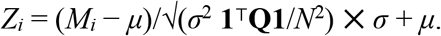

Voxelwise correction for false discovery rate (5% level) was performed using the two-stage linear step-up procedure ^36^.

The random-effects variance across teams was estimated using an analog to the tau-squared statistic used in meta-analysis. We used the following estimator to account for the interstudy correlation and provide an unbiased estimate of the between-team variance,

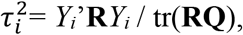

where *Y*_*i*_ is the vector of T statistics across teams at a given voxel *i*, **Q** is the correlation matrix across teams (pooling over all voxels), and **R** is the centering matrix (**R**= **I - 11**^T^/N); this is just the usual sample variance except N−1 is replaced by tr(**RQ**).

### Prediction markets

The second main goal of the Neuroimaging Analysis Replication and Prediction Study (NARPS) was to test the degree to which researchers in the field can predict results, using prediction markets ^5–7,37^. We invited team members (researchers that were members of one of the analysis teams) and non-team members (researchers that were neither members of any of the analysis teams nor members of the NARPS research group) to participate in a prediction market ^5,38^ to measure peer beliefs about the fraction of teams reporting significant whole-brain corrected results for each of the nine hypotheses. The prediction markets were conducted 1.5 months after all teams had submitted their analysis of the fMRI dataset. Thus, team members had information about the results of their specific team, but not about the results of any other team.

Similar to previous studies^5–7,20^, participants in the prediction markets were provided with monetary endowments (100 Tokens, worth $50) and traded on the outcome of the hypotheses via a dedicated online market platform (see Supplementary Figure 12). Each hypothesis constitutes one asset in the market, with asset prices predicting the fraction of teams reporting significant whole-brain corrected results for the corresponding ex-ante hypothesis examined by the analysis teams using the same dataset. Trading on the prediction markets was incentivized, i.e., traders were paid based on their performance in the markets.

#### Recruitment

For the “non-team members” prediction market, we invited participants via social media (mainly Facebook and Twitter) and e-mails. The invitation contained a link to an online form on the NARPS website (www.narps.info) where participants could sign up using their email address.

Participants for the “team members” prediction market were invited via email, after all teams submitted their results, directing them to an independent registration form (with identical form fields) to separate participants for the two prediction markets already at the time of registration. Note that team members initially were not aware that they would be invited to participate in a separate prediction market after they had analyzed the data. The decision to implement a second market, consisting of traders with partial information about the fundamental values (i.e., the team members) was made after the teams got access to the fMRI dataset. Thus, team members were only invited to participate in the market after all teams had submitted their analysis results. Once the registration for participating in the prediction markets had been closed, we reconciled the sign-ups with the list of team members to ensure that team members did not mistakenly end up in the “non-team members” prediction market and vice versa.

In addition to their email addresses, which were used as the only key to match registrations, accounts in the market platform, and the teams’ analysis results, registrants were required to provide the following information during sign-up: *(i)* name, *(ii)* affiliation, *(iii)* position (PhD candidate, Post-doctoral researcher, Assistant Professor, Senior Lecturer, Associate Professor, Full Professor, Other), *(iv)* years since PhD, *(v)* gender, *(vi)* age, *(vii)* country of residence, *(viii)* self-assessed expertise in neuroimaging (Likert scale ranging from 1 to 10), *(ix)* self-assessed expertise in decision sciences (Likert scale ranging from 1 to 10), *(x)* preferred mode of payment (Amazon.de voucher, Amazon.com voucher, PayPal payment), and *(xi)* whether they are a team member of any analysis team (yes / no). The invitations to participate in the prediction markets were first distributed on April 9, 2019; the registration closed on April 29, at 4pm UTC. Upon close of the registration, all participants received a personalized email containing a link to the web-based market software and their login-credentials. The prediction markets opened on May 2, 2019 at 4pm UTC and closed on May 12, 2019 at 4pm UTC.

#### Information available to participants

All participants had access to detailed information about the data collection, the experimental protocol, the ex-ante hypotheses, the instructions given to the analysis teams, references to related papers, and detailed instructions about the prediction markets via the NARPS website (www.narps.info).

#### Implementation of prediction markets

To implement the prediction markets, we used a newly developed web-based framework dedicated for conducting continuous-time online market experiments, inspired by the trading platform in the Experimental Economics Replication Project (EERP^7^) and the Social Sciences Replication Project (SSRP^6^). Similar to these previous implementations, there were two main views on the platform: (i) the market overview and (ii) the trading interface. The market overview showed the nine assets (i.e., one corresponding to each hypothesis) in tabular format, including information on the (approximate) current price for buying a share and the number of shares held (separated for long and short positions) for each of the nine hypotheses. Via the trading interface, which was shown after clicking on any of the hypotheses, the participant could make investment decisions and view price developments for the particular asset (see Supplementary Figure 12).

Note that initially, there was an error in the labelling of two assets (i.e., hypotheses) in the trading interface and the overview table of the web-based trading platform (the more detailed hypothesis description available via the info symbol on the right hand side of the overview table contained the correct information): Hypotheses 7 and 8 mistakenly referred to negative rather than positive effects of losses in the Amygdala. One of the participants informed us about the inconsistency between the information on the trading interface and the information provided on the website on May 6. The error was corrected immediately on the same day and all participants were informed about the mistake on our part via a personal email notification (on May 6, 2019, 3:28pm UTC), pointing out explicitly which information was affected and asking them to double-check their holdings in the two assets to make sure that they are invested in the intended direction.

#### Trading and market pricing

In both prediction markets, traders were endowed with 100 Tokens (the experimental currency unit). Once the markets opened, these Tokens could be used to trade shares in the assets (i.e., hypotheses). Unlike prediction markets on binary outcomes (e.g., the outcomes of replications as in previous studies^6,7^), for which market prices were typically interpreted as the predicted probability of the outcome to occur ^39^(though see^40^ and^41^ for caveats), the prediction markets accompanying the team analyses in the current study were implemented in terms of vote-share-markets. Hence, the prediction market prices serve as measures of the aggregate beliefs of traders for the fraction of teams reporting that the hypotheses were supported and can fluctuate between 0 (no team reported a significant result) and 1 (all teams reported a significant result).

Prices were determined by an automated market maker implementing a logarithmic market scoring rule^42^. At the beginning of the markets, all assets were valued at a price of 0.50 Tokens per share. The market maker calculated the price of a share for each infinitesimal transaction and updated the price based on the scoring rule. This ensured both that trades were always possible even when there was no other participant with whom to trade and that participants had incentives to invest according to their beliefs^43^. The logarithmic scoring rule uses the net sales (shares held - shares borrowed) the market maker has done so far in a market to determine the price for an infinitesimal trade as *p* = *e*^*s/b*^ / (*e*^*s/b*^ + 1). The parameter *b* determines the liquidity provided by the market maker and controls how strongly the market price is affected by a trade. We set the liquidity parameter to *b* = 100, implying that by investing 10 Tokens, traders could move the price of a single asset from 0.50 to about 0.55.

Investment decisions for a particular hypothesis were made from the market’s trading interface. In the trading overview, participants could see the (approximate) price of a new share, the number of shares they currently held (separated for long and short positions), and the number of Tokens their current position was worth if they liquidated their shares. The trading page also contained a graph depicting previous price developments. To make an adjustment to their current position, participants could choose either to increase or decrease their position by a number of Tokens of their choice. Supplementary Figure 12 depicts screenshots of the web-based software implementation. The trading procedures and market pricing are described in more detail in Camerer et al.^7^.

#### Incentivization

Once the markets had been closed, the true “fundamental value” (FV) for each asset (i.e., the fraction of teams that reported a significant result for the particular hypothesis) was determined and gains and losses were calculated as follows: If holdings in a particular asset were positive (i.e., the trader acted as a net buyer), the payout was calculated as the fraction of analysis teams reporting a significant result for the associated hypothesis multiplied by the number of shares held in the particular asset; If a trader’s holdings were negative (i.e., the trader acted as a net seller), the (absolute) amount of shares held was valued at the price differential between 1 and the fraction of teams reporting a significant result for the associated hypothesis.

Any Tokens that had not been invested into shares when the market closed were voided. Any Tokens awarded as a result of holding shares were converted to USD at a rate of 1 Token = $0.5. The final payments were transferred to participants during the months May to September 2019 in form of Amazon.com giftcards, Amazon.de giftcards, or PayPal payments, depending on the preferred mode of payment indicated by the participants upon registration for participating in the prediction markets.

#### Participants

In total, 96 “team members” and 91 “non-team members” signed up to participate in the prediction markets. N = 83 “team members” and N = 65 “non-team members” actively participated in the markets. The number of traders active in each of the assets (i.e., hypotheses) ranged from 46 to 76 (*m* = 56.4, *sd* = 8.9) in the “team members” set of markets and from 35 to 58 (*m* = 47.1, *sd* = 7.9) in the “non-team members” set of markets. See Supplementary Table 11 for data about trading volume on the prediction markets.

Of the participants, 10.2% did not work in academia (but hold a PhD), 34.2% were PhD students, 43.3% were post-docs or assistant professors, 7.5% were lecturers or associate professors, and 4.8% were full professors. 27.8% of the participants were female. The average time spent in academia after obtaining the PhD was 4.1 years. The majority of the participants resided in Europe (46.3%) and North America (46.3%).

#### Pre-Registration

All analyses of the prediction markets data reported were pre-registered at https://osf.io/pqeb6/. The pre-registration was completed after the markets opened, but before the markets closed. Only one member of the NARPS research group, Felix Holzmeister, had any information about the prediction market prices before the markets closed (as he monitored the prediction markets). He was not involved in writing the pre-registration. Only two members of the NARPS research group, Rotem Botvinik-Nezer and Tom Schonberg, had any information about the results reported by the 70 analyses teams before the prediction markets closed. Neither of them were involved in writing the pre-registration either.

For additional details on the prediction markets, see the Supplementary Materials.

## Acknowledgements

Neuroimaging data collection, performed at Tel Aviv University, was supported by the Austrian Science Fund (P29362-G27), the Israel Science Foundation (ISF, 1798/15 and 2004/15 granted to Tom Schonberg) and the Swedish Foundation for Humanities and Social Sciences (NHS14- 1719:1). Hosting of the data on OpenNeuro supported by NIH grant R24MH117179. Thanks to Michael C. Frank, Yaniv Assaf and Nathaniel Daw for helpful comments on an earlier draft. Thanks to the Texas Advanced Computing Center for providing computing resources for preprocessing of the data, and the Stanford Research Computing Facility for hosting the data. D. Wisniewski was supported by the Research Foundation Flanders (FWO, fwo.be), and the European Union’s Horizon 2020 research and innovation programme (https://ec.europa.eu/programmes/horizon2020/en) under the Marie Skłodowska-Curie grant agreement No 665501. L. Tisdall acknowledges the University of Basel Research Fund for Junior Researchers. C.B. Calderon was supported by grant 12O7719N from the Science Foundation Flanders. E. Lesage was supported by grant 12T2517N from the Science Foundation Flanders. A. Eed was supported by a predoctoral fellowship La Caixa-Severo Ochoa from Obra Social La Caixa and also acknowledges Comunidad de Cálculo Científico del CSIC for the HPC usage. C. Lamm was supported by the Viennese Science and Technology Fund (WWTF VRG13-007) and Austrian Science Fund (FWF P 32686). A. Losecaat Vermeer was supported by the Viennese Science and Technology Fund (WWTF VRG13-007). L. Zhang was supported by the National Natural Science Foundation of China (No. 71801110), MOE (Ministry of Education in China) Project of Humanities and Social Sciences (No. 18YJC630268), and China Postdoctoral Science Foundation (No. 2018M633270).

## Author contributions

- NARPS management team: R. Botvinik-Nezer, F. Holzmeister, C.F. Camerer, A. Dreber, J. Huber, M. Johannesson, M. Kirchler, R.A. Poldrack and T. Schonberg.
- fMRI dataset- experiment design: R. Iwanir, J. Durnez, R.A. Poldrack and T. Schonberg.
- fMRI dataset- data collection: R. Iwanir and T. Schonberg.
- fMRI dataset- preprocessing, quality assurance and data sharing: R. Botvinik-Nezer, K. Gorgolewski, R.A. Poldrack and T. Schonberg.
- Analysis teams- recruitment, point of contact and management: R. Botvinik-Nezer, R.A. Poldrack and T. Schonberg.
- Analysis teams- analysis of the submitted results and statistical maps: R.A. Poldrack, T.E. Nichols, J.A. Mumford, J-.B. Poline, A. Perez, R. Botvinik-Nezer, and T. Schonberg.
- Code review: T. Glatard. and K. Dadi.
- Prediction markets- design and management: F. Holzmeister, C.F. Camerer, A. Dreber, J. Huber, M. Johannesson and M. Kirchler
- Prediction markets- analysis: F. Holzmeister, R. Botvinik-Nezer, C.F. Camerer, A. Dreber, J. Huber, M. Johannesson, M. Kirchler, S. Kupek, R.A. Poldrack and T. Schonberg.
- Writing the manuscript: R. Botvinik-Nezer, F. Holzmeister, A. Dreber, J. Huber, M. Johannesson, M. Kirchler, T.E. Nichols, R.A. Poldrack and T. Schonberg.
- The rest of the authors participated as members of analysis teams and reviewed and edited the manuscript.

