## Supplementary Materials for "Variability in the analysis of a single neuroimaging dataset by many teams"

Rotem Botvinik-Nezer[1,2], Felix Holzmeister[3], Colin F. Camerer[4], Anna Dreber[3,5], Juergen Huber[3], Magnus Johannesson[5], Michael Kirchler[3], Roni Iwanir[1], Jeanette A. Mumford[6], Alison Adcock[7], Paolo Avesani[8,9], Blazej Baczkowski[10], Aahana Bajracharya[11], Leah Bakst[12], Sheryl Ball[13], Marco Barilari[14], Nadège Bault[9], Derek Beaton[15], Julia Beitner[16,17], Roland Benoit[10], Ruud Berkers[10], Jamil Bhanji[18], Bharat Biswal[19,20], Sebastian Bobadilla-Suarez[21], Tiago Bortolini[22], Katherine Bottenhorn[23], Alexander Bowring[24], Senne Braem[25], Hayley Brooks[26], Emily Brudner[18], Cristian Calderon[27], Julia Camilleri[28,29], Jaime Castrellon[7], Luca Cecchetti[30], Edna Cieslik[29,31], Zachary Cole[32], Olivier Collignon[14], Robert Cox[33], William Cunningham[34], Stefan Czoschke[17], Kamalaker Dadi[35], Charles Davis[36], Alberto De Luca[37], Mauricio Delgado[18], Lysia Demetriou[38,39], Jeffrey Dennison[40], Xin Di[19,41], Erin Dickie[34,42], Ekaterina Dobryakova[43], Claire Donnat[44], Juergen Dukart[29,31], Niall W. Duncan[45,46], Joke Durnez[47], Amr Eed[48], Simon Eickhoff[29,31], Andrew Erhart[26], Laura Fontanesi[49], G. Matthew Fricke[50], Adriana Galvan[51], Remi Gau[52], Sarah Genon[31], Tristan Glatard[53], Enrico Glerean[54], Jelle Goeman[55], Sergej Golowin[45,46], Carlos González-García[27], Krzysztof Gorgolewski[44], Cheryl Grady[15,34], Mikella Green[7], João Guassi Moreira[51], Olivia Guest[21], Shabnam Hakimi[7], J. Paul Hamilton[56], Roeland Hancock[36], Giacomo Handjaras[30], Bronson Harry[57], Colin Hawco[42,58], Peer Herholz[59], Gabrielle Herman[42], Stephan Heunis[60,61], Felix Hoffstaedter[28,29], Jeremy Hogeveen[62], Susan Holmes[44], Chuan-Peng Hu[63], Scott Huettel[7], Matthew Hughes[64], Vittorio Iacovella[65], Alexandru Iordan[66], Peder Isager[60], Ayse Ilkay Isik[67], Andrew Jahn[66], Matthew Johnson[32], Tom Johnstone[64], Michael Joseph[42], Anthony Juliano[43,68], Joseph Kable[69], Michalis Kassinopoulos[59], Cemal Koba[30], Xiang-Zhen Kong[70], Timothy Koscik[71], Nuri Erkut Kucukboyaci[43,72], Brice Kuhl[73], Sebastian Kupek[3], Angela Laird[23], Claus Lamm[74], Robert Langner[29,31], Nina Lauharatanahirun[69,75], Hongmi Lee[76], Sangil Lee[69], Alexander Leemans[37], Andrea Leo[30], Elise Lesage[27], Flora Li[77,78], Monica Li[36], Phui Cheng Lim[32], Evan Lintz[32], Schuyler Liphardt[62], Annabel Losecaat Vermeer[74], Bradley Love[21], Michael Mack[34], Norberto Malpica[79], Theo Marins[22], Camille Maumet[80], Kelsey McDonald[7], Joseph McGuire[12], Helena Melero[79], Adriana Méndez Leal[51], Benjamin Meyer[63], Kristin Meyer[81], Paul Mihai[10,82], Georgios Mitsis[59], Jorge Moll[22], Dylan Nielson[33], Gustav Nilsonne[83,84], Michael Notter[85], Emanuele Olivetti[8,9], Adrian Onicas[30], Paolo Papale[30], Kaustubh Patil[28,29], Jonathan E. Peelle[11], Alexandre Pérez[86], Doris Pischedda[9,87], Jean-Baptiste Poline[59,88], Yanina Prystauka[36,89], Shruti Ray[19], Patricia Reuter-Lorenz[66], Richard Reynolds[33], Emiliano Ricciardi[30], Jenny Rieck[15], Anais Rodriguez-Thompson[81], Anthony Romyn[34], Taylor Salo[23], Gregory Samanez-Larkin[7], Emilio Sanz-Morales[90], Margaret Schlichting[34], Douglas Schultz[32], Qiang Shen[91], Margaret Sheridan[81], Fu Shiguang[91], Jennifer Silvers[51], Kenny Skagerlund[56], Alec Smith[13], David Smith[40], Peter Sokol-Hessner[26], Simon Steinkamp[31], Sarah Tashjian[51], Bertrand Thirion[35,92], John Thorp[93], Gustav Tinghög[56], Loreen Tisdall[49], Steven Tompson[75], Claudio Toro-Serey[12], Juan Torre[35,92], Leonardo Tozzi[44], Vuong Truong[45,46], Luca Turella[65], Anna E. van 't Veer[94], Tom Verguts[27], Jean Vettel[75,95], Sagana Vijayarajah[34], Khoi Vo[7], Matthew Wall[38,96], Wouter D. Weeda[94], Susanne Weis[29,97], David White[64], David Wisniewski[27], Alba Xifra-Porxas[59], Emily Yearling[36], Sangsuk Yoon[98], Rui Yuan[44], Kenneth Yuen[99], Lei Zhang[74], Xu Zhang[36], Joshua Zosky[32], Thomas E. Nichols[24,100,*], Russell A. Poldrack[44,*], Tom Schonberg[1,*]

### Analysis teams results

The Spearman correlation of the distance of the outcome from 0.5 (i.e., how consistent the results were across teams) and the mean confidence level across hypotheses was positive ($r = 0.69$, $p = 0.039$, $n = 70$), indicating that when variability of the outcome across teams was smaller, the teams were more confident in their results. The Spearman correlation between the distance of the outcome from 0.5 and the mean estimated similarity to other teams was not significant ($r = 0.40$, $p = 0.286$, $n = 70$).

### Variability of unthresholded statistical maps

No teams were consistently anticorrelated with the mean pattern across all hypotheses, though three teams showed a correlation of $r < 0.2$ with the mean pattern across hypotheses, whereas 32 teams showed correlations of $r > 0.7$ with the mean pattern.

### Prediction markets

A limitation of the prediction markets part of the study is that the number of observations for each set of prediction markets is low, as the number of observations for each set of prediction markets equals the number of hypotheses ($n = 9$) tested by the teams in the fMRI dataset. This meant that we had nine prediction market observations for "team members" and nine prediction market observations for "non-team members". These were aggregated market observations about predictions of the fraction of teams reporting significant results for each hypothesis (bounded between 0 and 1). The low number of observations implied that the statistical power to find statistically significant effects was limited, and the test results should therefore be interpreted cautiously.

#### Prediction markets results

**Traders self-ranked expertise.** On average, participants' self-reported expertise in neuroimaging (Likert scale from 1 to 10) was 6.54 ($sd = 1.93$) for the "team members" prediction market and 5.98 ($sd = 2.39$) for the "non-team members" prediction market, respectively (Welch two-sample $t$-test: $t(173.19) = 1.77$, $p = 0.078$). The mean self-reported expertise in decision sciences (Likert scale from 1 to 10) was significantly higher for the "non-team members" ($mean = 5.13$, $sd = 2.36$) compared to the "team members" ($mean = 4.23$, $sd = 2.46$) prediction market (Welch two-sample $t$-test: $t(184.97) = 2.56$, $p = 0.011$). These tests comparing the value of the variables between the two samples were not pre-registered and are included for descriptive purposes.

**Exploratory analyses.** Although not stated in the pre-analysis plan, we examined the correlation between participants' final payoffs, as an indicator of market performance and prediction accuracy, with participants' self-reported expertise in neuroimaging and decision sciences. The Spearman correlations between payoffs and self-rated expertise turn out to be low in magnitude and statistically insignificant for expertise in both neuroimaging ($r = 0.06$, $p = 0.45$, $n = 148$) and decision sciences ($r = -0.07$, $p = 0.369$, $n = 148$). This exploratory result also holds if we examine Spearman correlations for "team members" and "non-team members" separately (expertise in neuroimaging: "non-team members", $r = 0.19$, $p = 0.141$, $n = 65$; "team-members", $r = -0.12$, $p = 0.273$, $n = 83$; expertise in decision sciences: "non-team members", $r = 0.04$, $p = 0.745$, $n = 65$; "team-members", $r = 0.02$, $p = 0.829$, $n = 83$).

To explore whether and how market prices (i.e., market's predictions) aggregate traders' private information over time, we calculated the absolute error of the market price from the fundamental value on an hourly basis (average price of all transactions within an hour), resulting in a time series of 240 observations (10 days x 24 hours; see Supplementary Figure 13). We ran two panel regressions with 18 cross-sections (i.e., nine hypotheses run for both sets of markets) and 240 time observations each. In the model (1), we regressed the absolute error on a binary prediction market indicator "team members" and control for linear time effect. The statistically significant coefficient for the team membership dummy ($\beta = -0.22$, $p < 0.001$) indicated that, on average, predictions in the "team members" prediction market were closer to the fundamental value than aggregate market's predictions in the "non-team members" prediction market. The positive coefficient for the time trend ($\beta = 4.41 \times 10^{-4}$, $p < 0.001$) in the model suggested that information aggregation got worse over time, i.e. that prices in both prediction markets tended to drift away from the fundamental value as time progressed. Adding the interaction term of the time trend and the prediction market indicator variable in model (2) revealed that prediction errors over time increased at a significantly higher rate in the "team members" prediction market compared to the "non-team members" prediction market. Despite the lower prediction errors in the "team members" prediction market, this suggests that information aggregation over time was more effective in the "non-team members" prediction market. The results are presented in Supplementary Table 12.

Concerning individual traders and how their opinions were incorporated in the market's predictions, we carried out two analyses for the "team members" prediction market only. First, Spearman correlations between the results their team has reported (a binary outcome) and their individual final holdings in the asset for each of the nine hypotheses range from 0.23 to 0.74 (all correlations are statistically significant, except for Hypothesis #7: $\rho_s = 0.23$, $p = 0.104$; for details, see Supplementary Table 9). In a second analysis, we calculated the percentage of trades in the "team members" prediction markets which are consistent with the results their team reported (i.e., whether they buy when their team reported a significant result in the hypothesized direction, but the market prices reflect "no significant result" and vice versa) for each of the nine hypotheses. The fractions of consistent trades ranged from 0.68 to 0.89. One-sample Wilcoxon signed-rank tests for a share of 0.5 revealed that the share of consistent trades was significantly higher than 50% ($z$-values range from 2.78 to 6.81; $p < 0.004$ for all tests; see Supplementary Table 9 for details). However, it turns out that inconsistent trades are disproportionately larger (in terms of volume) than consistent trades, explaining the systematic overvaluation of fundamental values.

In order to test whether overoptimism of traders in the team prediction market was the result of over-representation of teams reporting significant results, we computed the fraction of active traders that reported a significant result for each hypothesis. Overall, active traders in the teams prediction market were representative with respect to the overall results. The absolute differences in the fraction of significant results for active traders compared to all teams are small and vary from 0.021 to 0.088. For all hypotheses, the fraction of significant results for active traders lies within the 95% confidence intervals associated with the fraction of significant results reported by all teams, indicating that the active traders' information in the market are representative for the overall results. Moreover, for all hypotheses but one (Hypothesis #5), the fraction of significant results was lower for the active traders compared to all teams (see Supplementary Figure 11). Therefore, overoptimism of the traders in the teams prediction market could not be attributed to a biased outcome for these researchers.

### Supplementary Discussion

#### Analytic variability and its related factors

In NARPS, 70 analysis teams independently analyzed the same fMRI dataset to test the same nine ex-ante hypotheses which were based on the relevant scientific literature. Reported analysis outcomes demonstrated substantial variability in results across analysis teams. We further found that while the agreement between thresholded statistical maps was largely limited to regions with no active voxels, correlations between the unthresholded statistical maps across teams were moderate. Our exploratory analysis pointed out specific factors that significantly contributed to the variability. Higher estimated smoothness of the unthresholded statistical map, analyzing the data with FSL and using parametric correction methods were all related to more significant results. While the analysis software and correction method used are analytic choices directly made by each team, the estimated smoothness is a feature of the map and is affected by multiple earlier analytic choices. For example, exploratory analysis showed that modeling head movement was related to reduced estimated smoothness.

We did not find significant differences in results between analysis teams that chose to use the preprocessed (with fMRIPrep) shared dataset versus the teams that chose to use the raw dataset and preprocess the data by themselves. However, it should be noted that preprocessing includes many analytical procedures, and the effect of each specific procedure on the variability of final results was not directly tested here due to lack of power resulting from the multiple available options for each step.

The indications that correlated unthresolded statistical maps resulted in substantially different binary results across analysis teams suggested that a main source of the variability comes from the final stages of analysis: thresholding, correcting for multiple comparisons and anatomical ROI specifications. Although the general correction method used (parametric versus nonparametric) was found to be related to the final results, exploratory analysis applying a fixed threshold, correction method and anatomical ROI specification did not yield qualitatively more similar binary results compared to the reported ones (Supplementary Figure 9 and Supplementary Table 7). Nonetheless, correlated statistical maps should not necessarily produce similar binary results when applying the same threshold, since the correlation coefficient is not sensitive to overall scaling and thus correlated values could differ substantially in magnitude. Use of consistent thresholding and meta-analytic approaches provide another view on the heterogeneity (Supplementary Table 7). Hypotheses 2, 4, 5 & 6 all had at least 50% of teams showing activation on some thresholding approach and image-bsaed meta-analysis (IBMA) significance. While IBMA is based on the mean activation map, coordinate-based meta-analysis (CBMA) results can be driven by a subset of studies, and notably CMBA finds significance on all hypotheses except

#7; e.g. hypothesis 1 had over 50% activation and 1184 CBMA-significant voxels but none with IBMA, suggesting particular heterogeneity for that hypothesis' results.

There are several important analytic choices that could not be directly tested here. For example, as each hypothesis was related to a specific brain region, each team was required to choose an operative definition of the specific hypothesized region (i.e., in order to decide whether a significant activation was found within this region or not). Given the exact same thresholded statistical map, different teams could potentially conclude differentially [44]. Moreover, one of the three regions of interest in the current study was the ventromedial prefrontal cortex (vmPFC), for which there is no specific agreed-upon anatomical definition. This may have further contributed to variability across teams. However, we could not include this analytic choice in the tested model, as there were too many distinct methods used by the teams (e.g., different atlases, Neurosynth [45], visual examination, etc.) resulting in the lack of power to detect significant differences. Another important step we could not directly measure here was the general linear model specification. For example, modelling response time (RT) (or not) could potentially affect the results; the majority of teams (44) did not do so, but there were several different methods used by the teams that did. We did find several model specification errors that resulted in statistical maps that were anticorrelated with the majority of teams. While some of these errors might be related to the relative complexity of the particular task used here, other errors, such as those involving the inclusion of multiple correlated parameters in the model, likely generalize to all models.

It should also be noted that our results are conditional on the specific task we chose to use here, the mixed gambles task. This task is relatively complex, with multiple parametric modulators that could be (and were) modeled in a number of different ways. While this is a relatively representative task, a simpler task may have resulted in lower variance across pipelines (e.g., if there was less flexibility in the specification of the statistical model and region definitions).

**Prediction markets**

We used prediction markets to test the degree to which researchers from the field can predict the results. While traders in the "team members" prediction market had the data and knew their own results, traders from the "non-team members" reported significantly higher expertise in decision-sciences and are therefore assumed to be more familiar with the relevant literature. Nonetheless, we found that both groups of traders strongly overestimated the fraction of significant results. These results indicate that researchers in the field are over-optimistic with regard to the reproducibility of results across analysis teams. Nonetheless, team members predicted the relative plausibility of the hypotheses very well. Surprisingly, neither self-rated expertise in neuroimaging nor self-rated expertise in decision-sciences were related to better performance in the prediction markets (i.e., to better prediction of the results; see Supplementary Materials).

### Implications regarding previous findings with the mixed gamble task

There is a spectrum of concerns regarding the quality of research, ranging from replicability (the ability to reproduce a result in a new sample) to computational reproducibility (the ability to reproduce a result given data and analysis plans)[46]. Concerns over replicability across many areas of science have led to a number of projects in recent years that have attempted to assess the replicability of empirical findings across labs[6,20,37,47]. While such an undertaking would certainly be useful in the context of fMRI, the expense of fMRI data collection makes a large-scale replication attempt across many studies very unlikely. The present study does not broadly assess the replicability of neuroimaging research, but it does provide valuable insights, given that the design of the present study overlaps (in the equal indifference group) with the previous study of Tom et al.[8]. Out of the four primary claims made in the initial paper (reflecting significant outcomes on Hypotheses #1, #3 and #5, and a null outcome on Hypothesis #7), two were supported by a majority of teams in the present study. Moreover, as results largely differed for the equal indifference group (for which the design was similar to Tom et al.[8]) and the equal range group (for which the design was similar to De Martino et al[9].), mainly for the negative loss effect in the vmPFC (Hypothesis #5 vs. Hypothesis #6), inconsistent findings across these studies may be the result of the different designs they used. However, as the present study did not aim to directly test replicability of fMRI findings, but rather the variability across analysis pipelines, the implications are limited and should be interpreted with caution.

### General implications and proposed solutions

In this study, we assessed the degree to which results are reproducible across multiple analysts given a single dataset and pre-defined hypotheses. Our findings raise substantial concerns and indicate an urgent need for minimizing the effect of analytic choices on reported results[44]. Furthermore, our findings indicate that the further one gets from raw data the more divergent the results are. One implication of these findings is that meta-analyses should be more effective when using less processed data (i.e. unthresholded statistical maps versus thresholded statistical maps or activation coordinates) [cf. 48].

Importantly, the analysis teams who participated in the present study were not incentivized to find significant effects, which is thought to drive a number of questionable research practices (e.g., "p-hacking"[2]). Thus, the variability in the present results is more likely to reflect actual variability in the standard analytic methods used in the participating research groups and their interaction with the nature of the signals in the data, as well as model specification errors present for some teams. Moreover, the present study exclusively focused on univariate analysis. While this type of analysis is frequent, many fMRI studies in recent years have been using multivariate pattern analyses, which are less standardized and are therefore even more prone to be affected by specific analytic choices (although these methods may partially overcome the voxelwise thresholding stage). An open question is how the present results would generalize to those studies in which the researchers are motivated to detect a significant result (due to the prevalent bias for publication of significant results). We here note that our results imply substantial researcher degrees of freedom resulting in ample scope for p-hacking, as a significant result for each hypothesis could be reported based on at least four (based on the number of teams that reported a significant result for hypotheses 7-9) of the pipelines used in practice by analysis teams.

We propose that complex datasets should be analyzed using multiple analysis pipelines, preferably by more than one researcher, who would be blinded to the hypotheses of interest[23], and the results compared to ensure concordance across validated pipelines. The current study and future ones could point at the main analytic choices that lead to variable results. "Multiverse analysis" thus can be focused on those analytic choices to save required computational resources and allow a wider use across research groups. Previous studies in other fields have suggested different versions of "multiverse" analysis[26,27], but these have yet to be widely implemented. Meta-analysis methods can be used to draw conclusions based on multiple analysis pipelines and/or studies (when unthresholded statistical maps would be shared alongside neuroimaging publications). We believe this is a promising and important future direction, given the substantial influence of analytic choices on reported results. We also propose that the use of well-engineered and well-validated software tools instead of custom solutions, when appropriate, can help reduce the presence of errors and suboptimal analysis choices simply by the fact that these have been tested by multiple users and often employ more rigorous software engineering practices (but, importantly, should not be used as a "black box").

It is important to note, however, that concordance among different analysis pipelines does not necessarily imply that the conclusion of those analyses is correct. In the present study we do not have a "ground truth" regarding the effects (i.e., we do not know for certain whether each hypothesis is correct or not). Therefore, the present study provides crucial evidence and insights regarding the variability of results across analysis pipelines "in the wild" and its related factors, but not regarding the validity of each analytic choice or which analytic choices are the best ones. Future studies can use simulated data or null data, where the ground truth is known, to validate analysis workflows (e.g.[34,49]). These studies could potentially identify optimal analysis pipelines, on which the "multiverse analysis" could rely. We do not, however, believe that there is a single (or even a few) best analysis pipeline across studies[50,51]. Novel analysis methods are important for scientific discovery and progress, and different pipelines are optimal for different studies and scientific questions. Therefore, we suggest to focus on "multiverse analysis", while aggregating evidence across studies by sharing unthresholdd statistical maps and applying meta-analysis approaches. The discussed challenges and potential solutions are relevant far beyond neuroimaging, to any scientific field where the data are complex and there are multiple acceptable analysis workflows.

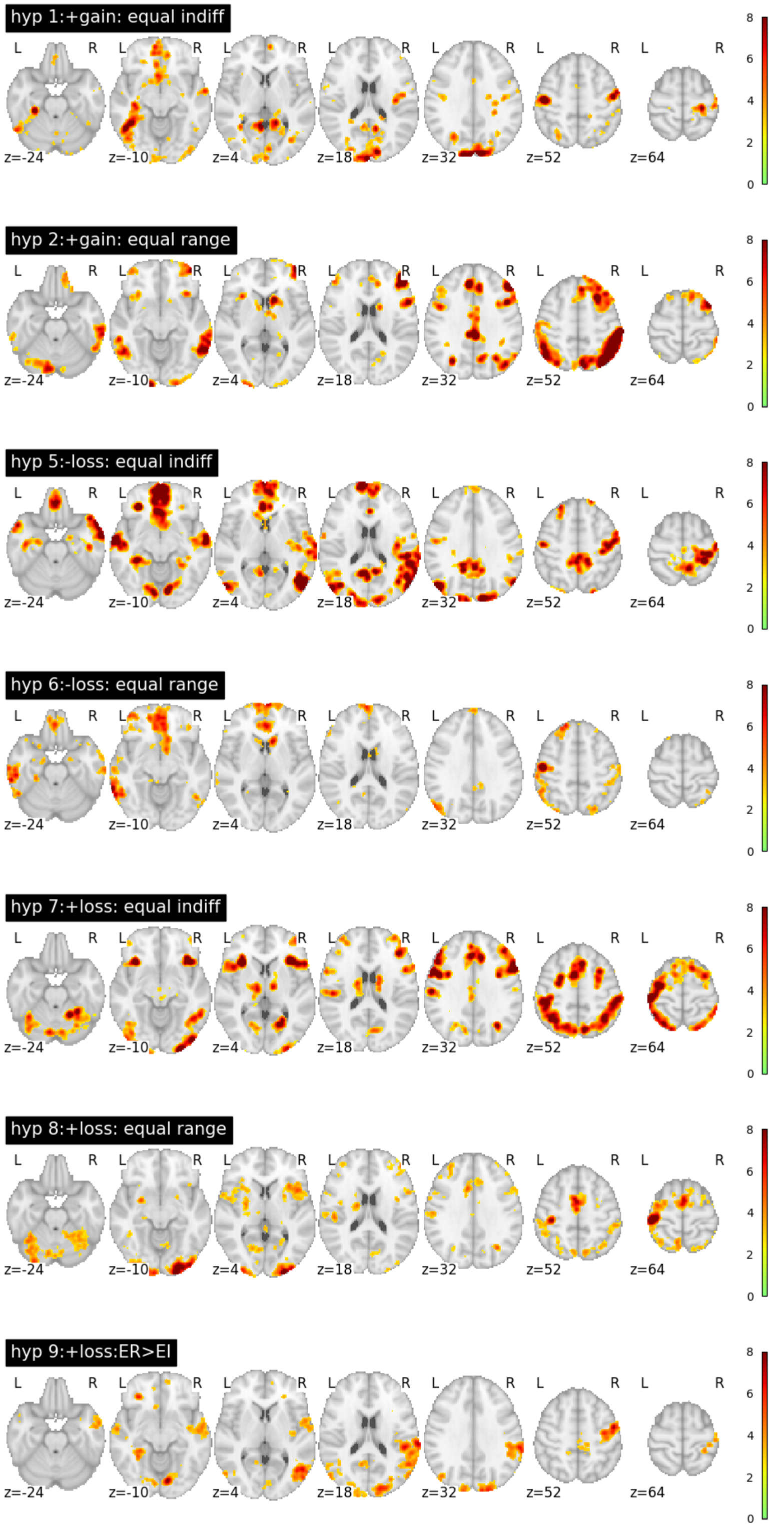


Supplementary Figure 1: Results from coordinate-based meta-analysis (CBMA) using activation likelihood estimation (ALE) across the thresholded statistical maps submitted by the analysis teams, separately for each hypothesis. Maps are thresholded at $p < 0.05$ corrected using false discovery rate. Images can be viewed at `https://identifiers.org/neurovault.collection:6049`.

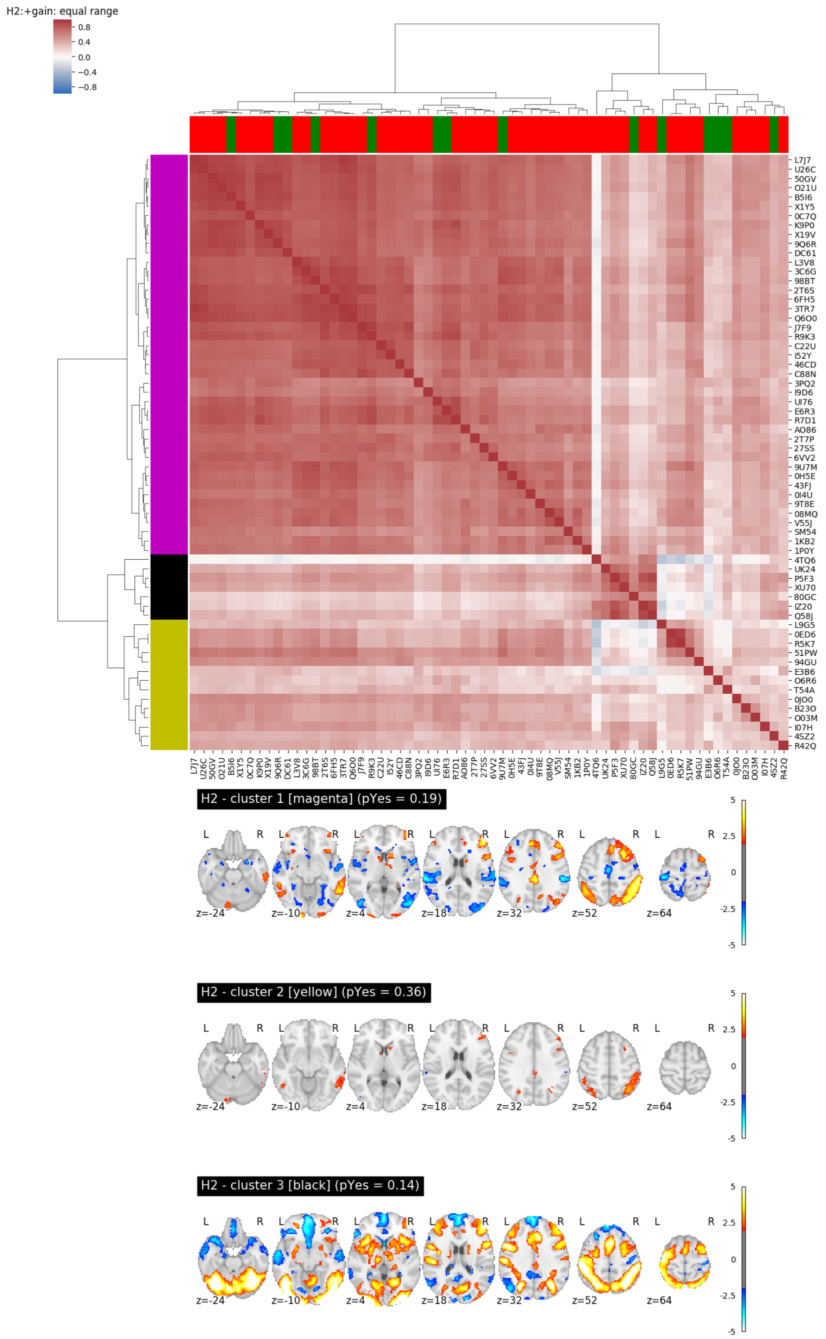


Supplementary Figure 2: Unthresholded map analysis for Hypotheses 2 and 4 (which both relate to the same contrast and group, but different regions). Top: Heatmap based on Spearman correlation between unthresholded statistical maps. Red / green color in the columns represent the decision regarding hypothesis 2 (no / yes, respectively). Bottom: Average of unthresholded images for each cluster (cluster colors in titles refer to colors in left margin of heatmap). Maps are thresholded at an uncorrected value of $Z > 2.0$ for visualization.

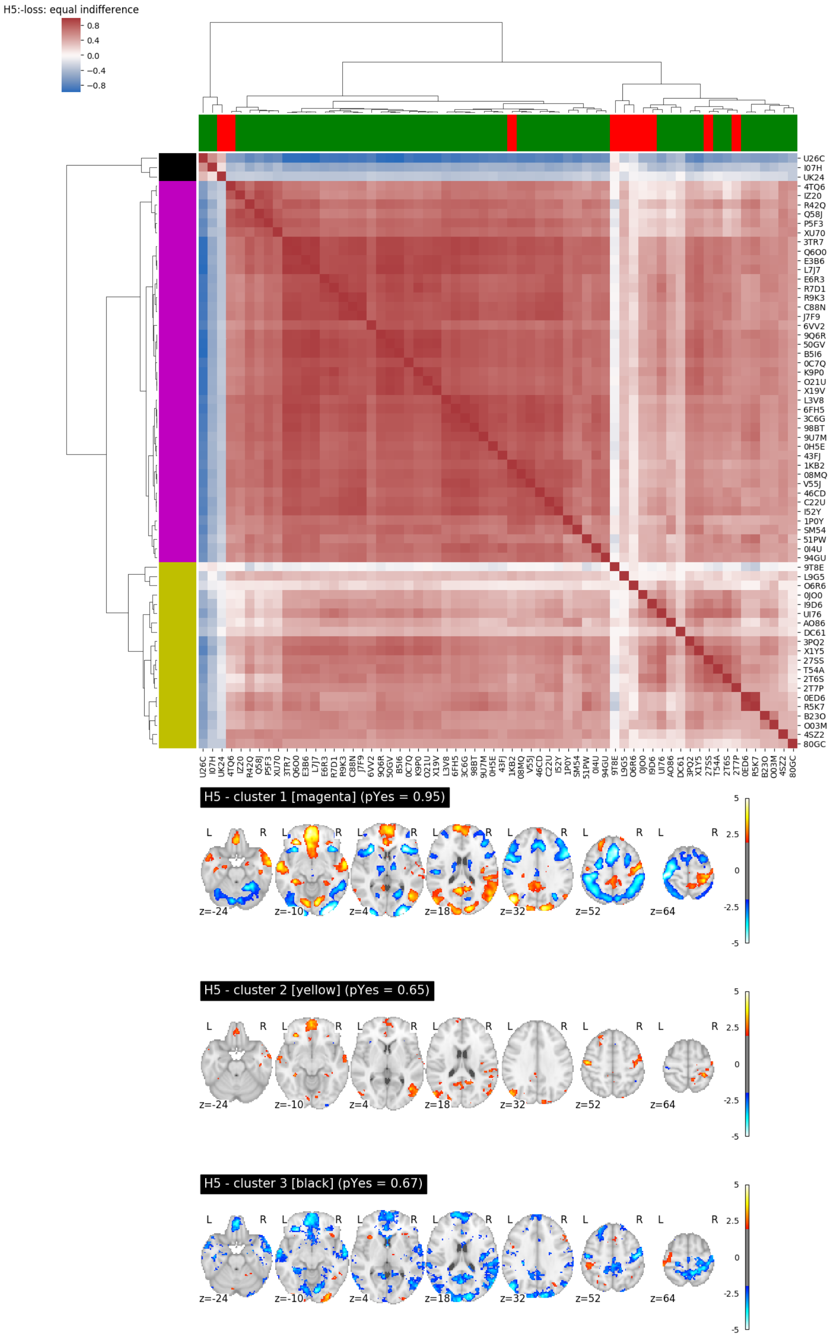


Supplementary Figure 3: Unthresholded map analysis for Hypothesis 5. Top: Heatmap based on Spearman correlation between unthresholded statistical maps. Red / green color in the columns represent the decision regarding hypothesis 5 (no / yes, respectively). Bottom: Average of unthresholded images for each cluster (cluster colors in titles refer to colors in left margin of heatmap). Maps are thresholded at an uncorrected value of $Z > 2.0$ for visualization.

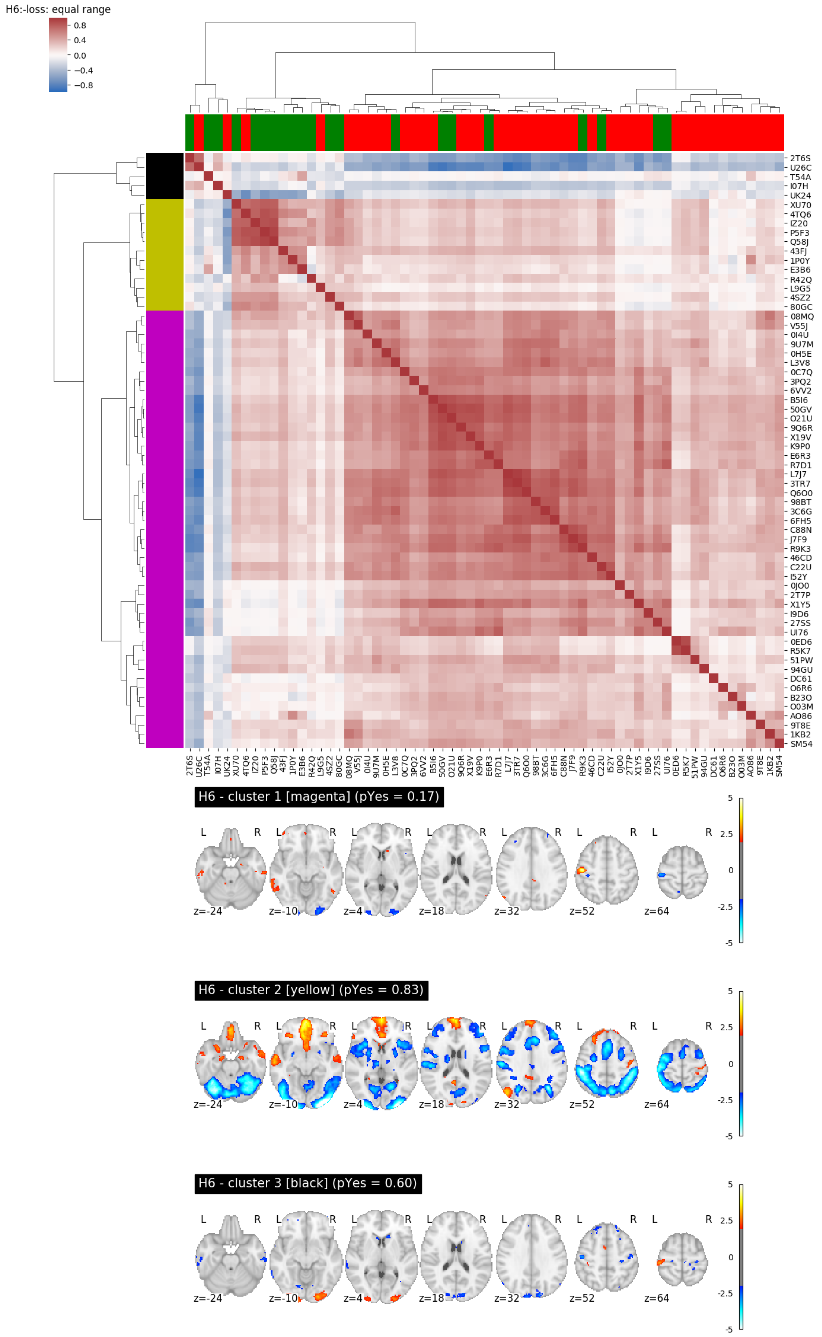


Supplementary Figure 4: Unthresholded map analysis for Hypothesis 6. Top: Heatmap based on Spearman correlation between unthresholded statistical maps. Red / green color in the columns represent the decision regarding hypothesis 6 (no / yes, respectively). Bottom: Average of unthresholded images for each cluster (cluster colors in titles refer to colors in left margin of heatmap). Maps are thresholded at an uncorrected value of $Z > 2.0$ for visualization.

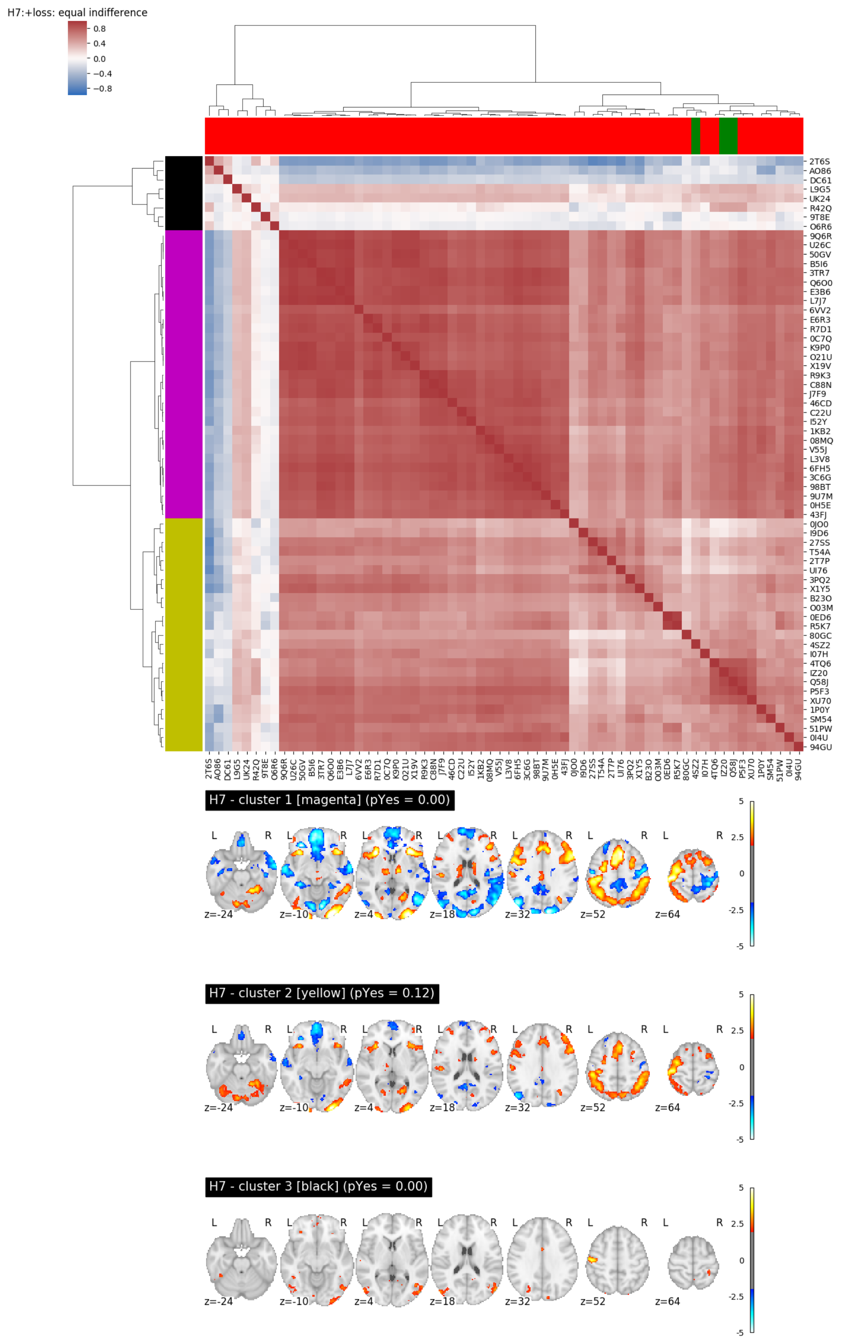


Supplementary Figure 5: Unthresholded map analysis for Hypothesis 7. Top: Heatmap based on Spearman correlation between unthresholded statistical maps. Red / green color in the columns represent the decision regarding hypothesis 7 (no / yes, respectively). Bottom: Average of unthresholded images for each cluster (cluster colors in titles refer to colors in left margin of heatmap). Maps are thresholded at an uncorrected value of $Z > 2.0$ for visualization.

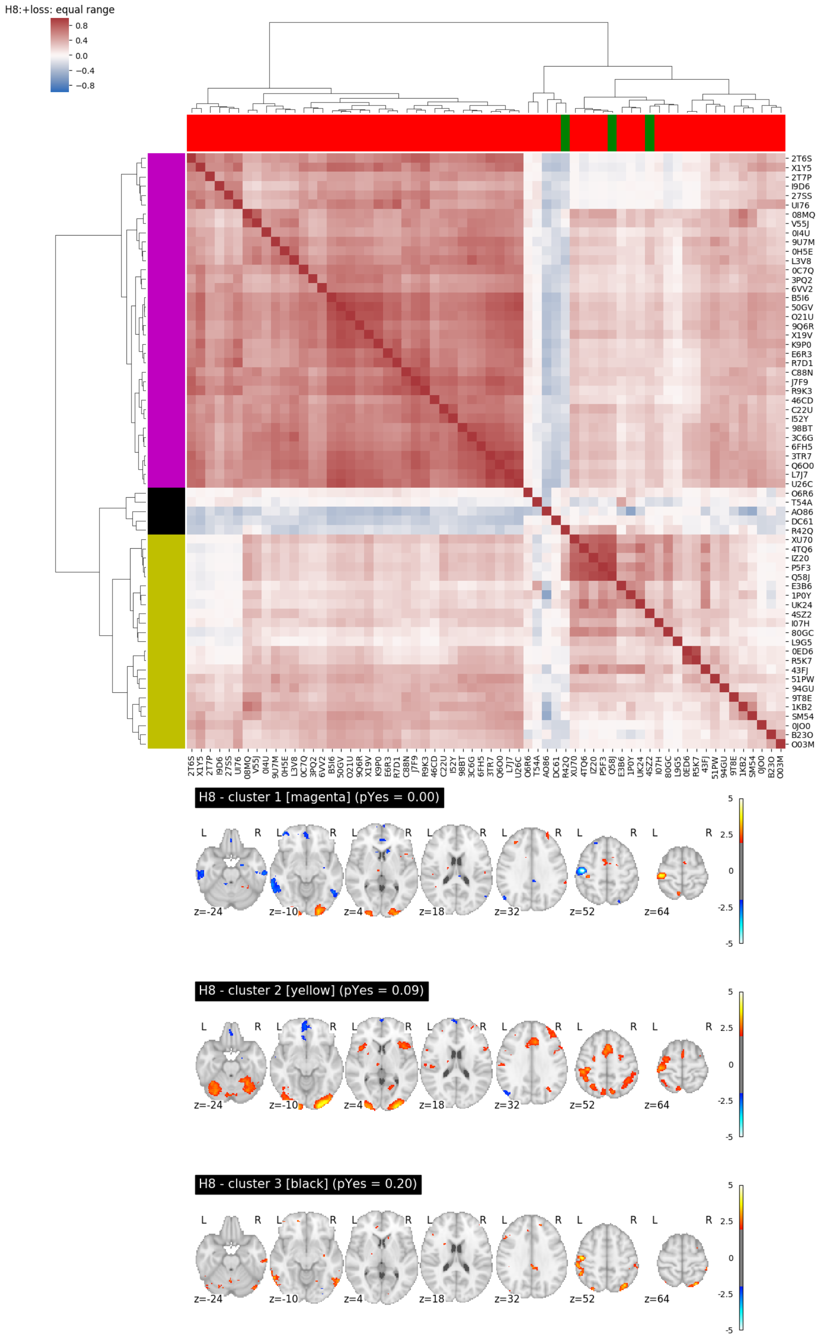


Supplementary Figure 6: Unthresholded map analysis for Hypothesis 8. Top: Heatmap based on Spearman correlation between unthresholded statistical maps. Red / green color in the columns represent the decision regarding hypothesis 8 (no / yes, respectively). Bottom: Average of unthresholded images for each cluster (cluster colors in titles refer to colors in left margin of heatmap). Maps are thresholded at an uncorrected value of $Z > 2.0$ for visualization.

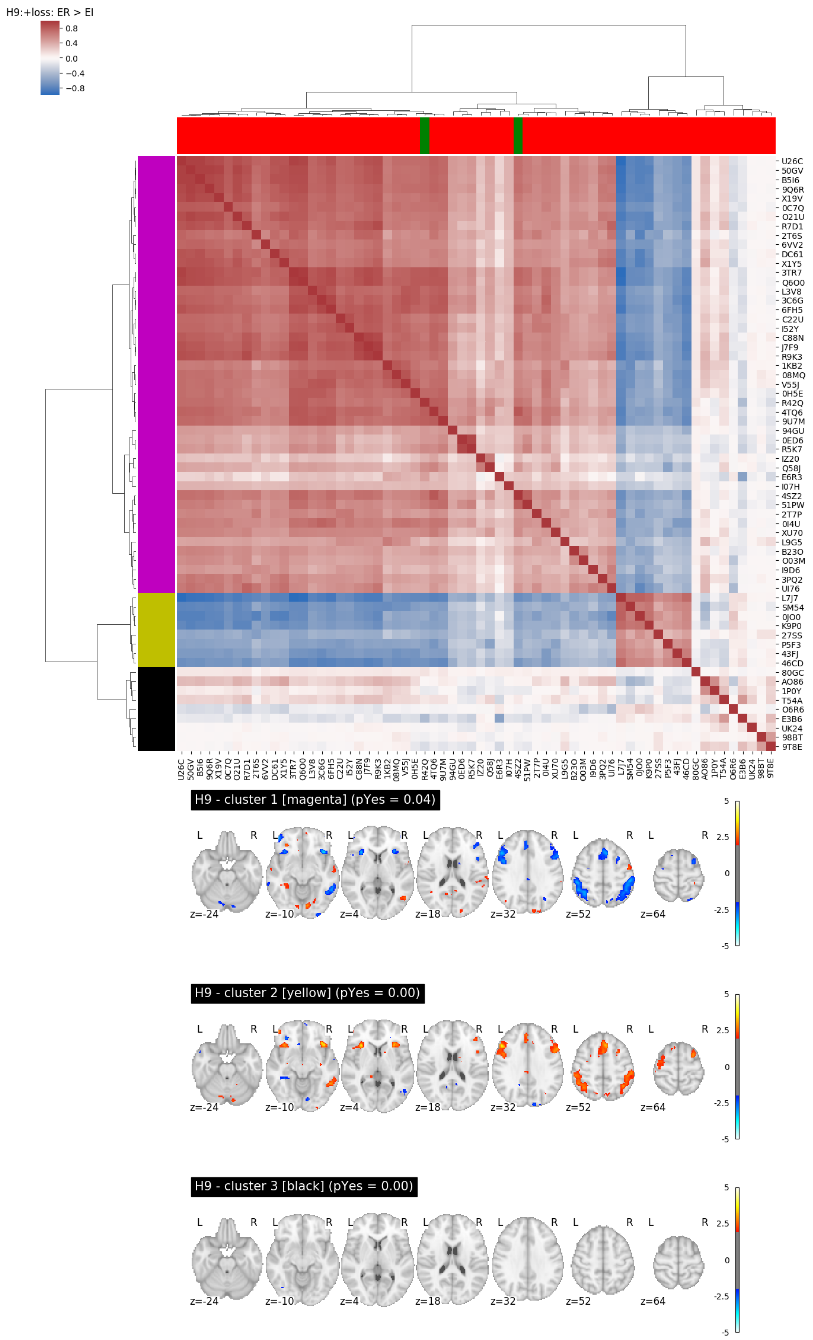

Supplementary Figure 7: Unthresholded map analysis for Hypothesis 9. Top: Heatmap based on Spearman correlation between unthresholded statistical maps. Red / green color in the columns represent the decision regarding hypothesis 9 (no / yes, respectively). Bottom: Average of unthresholded images for each cluster (cluster colors in titles refer to colors in left margin of heatmap). Maps are thresholded at an uncorrected value of $Z > 2.0$ for visualization.

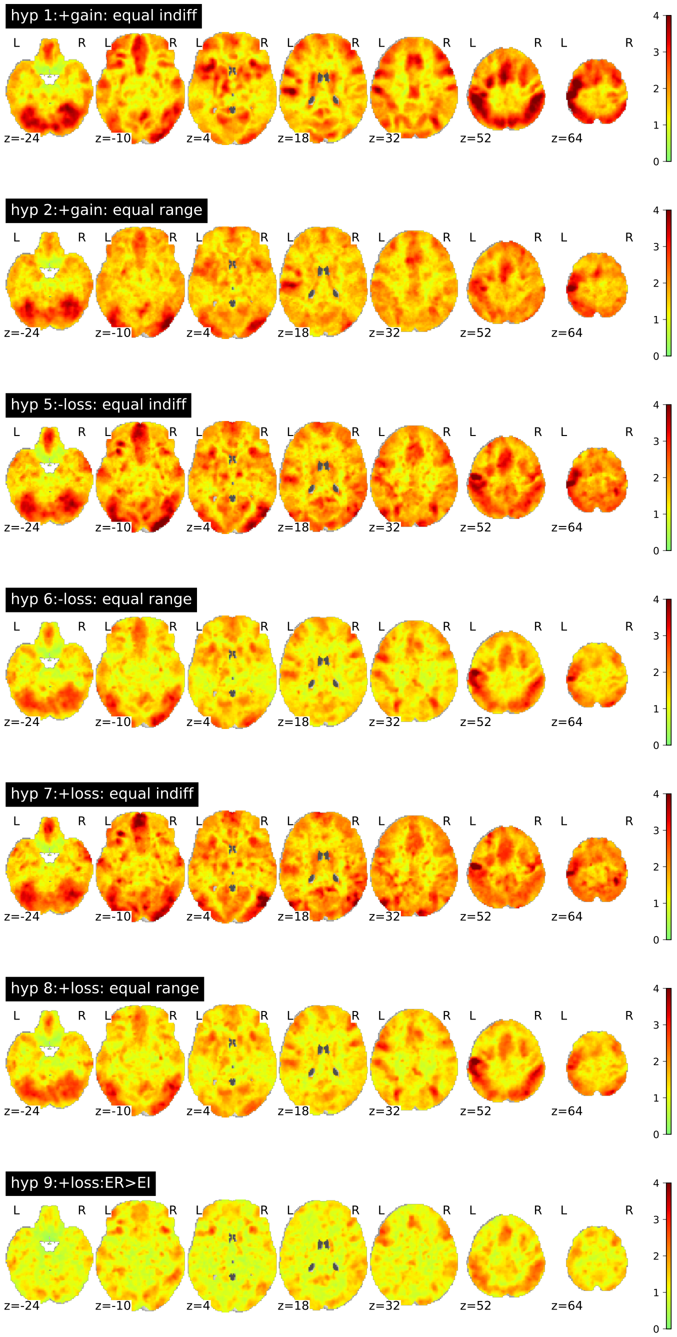


Supplementary Figure 8: Maps of estimated between-team variability (tau) at each voxel for each separate unthresholded map. Hypotheses #2 and #4 are not shown, as they share the same statistical maps as Hypothesis #1 and #3 respectively, which are for the same contrast and group but for different regions (see Table 1). Images can be viewed at `https://identifiers.org/neurovault.collection:6050`.

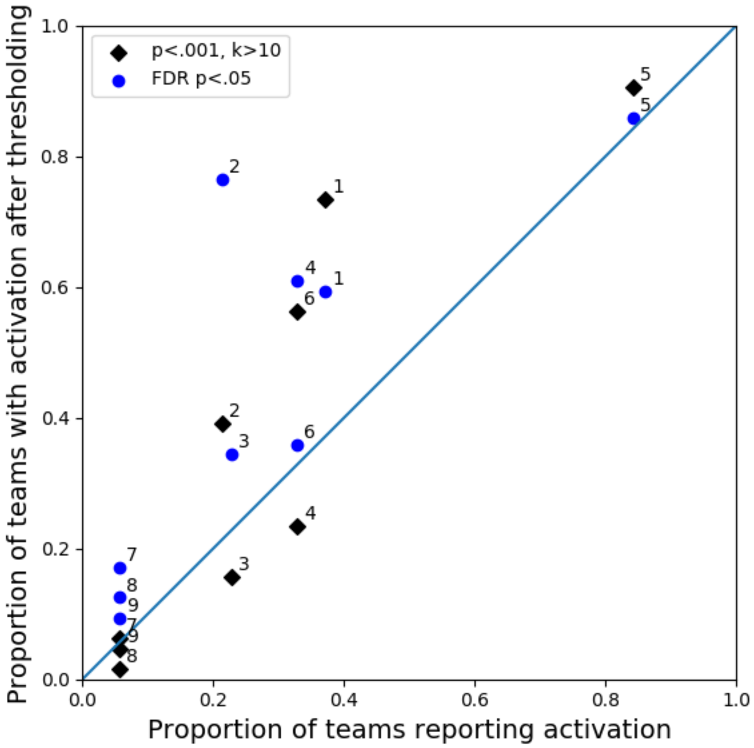


Supplementary Figure 9: Activation for each hypothesis as determined using consistent thresholding and ROI selection across teams (y-axis), versus proportion of teams reporting activation (x-axis). Numbers next to each symbol represent the hypothesis number for each point.

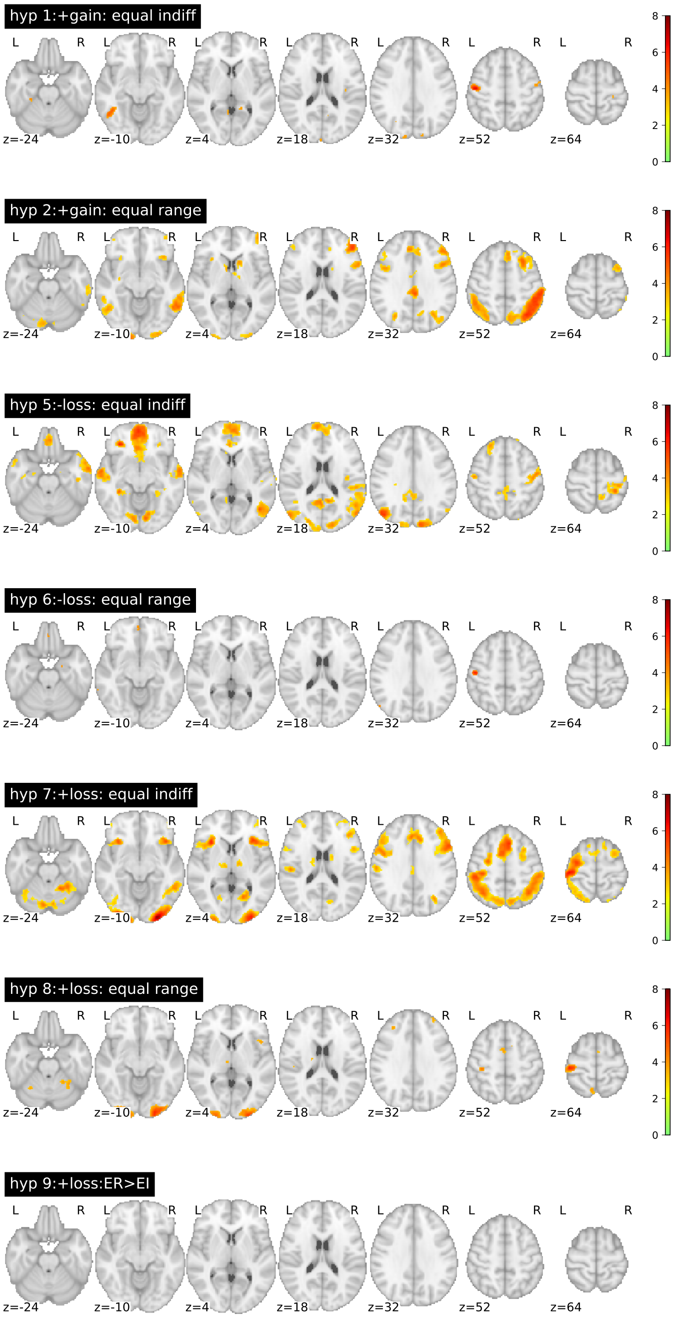


Supplementary Figure 10: Image-based meta-analysis (IBMA) results. A consensus analysis was performed on the unthresholded statistical maps submitted by the analysis teams to obtain a group statistical map for each hypothesis, accounting for the correlation between teams due to the same underlying data. Maps are presented for each hypothesis showing voxels (in color) where the group statistic was significantly greater than zero after voxelwise correction for false discovery rate ($p < .05$). Color bar reflects statistical value (Z) for the meta-analysis. Hypotheses #1 and #3, as well as hypotheses #2 and #4, share the same statistical maps as the hypotheses are for the same contrast and group, but for different regions (see Table 1). Images can be viewed at `https://identifiers.org/neurovault.collection:6051`.

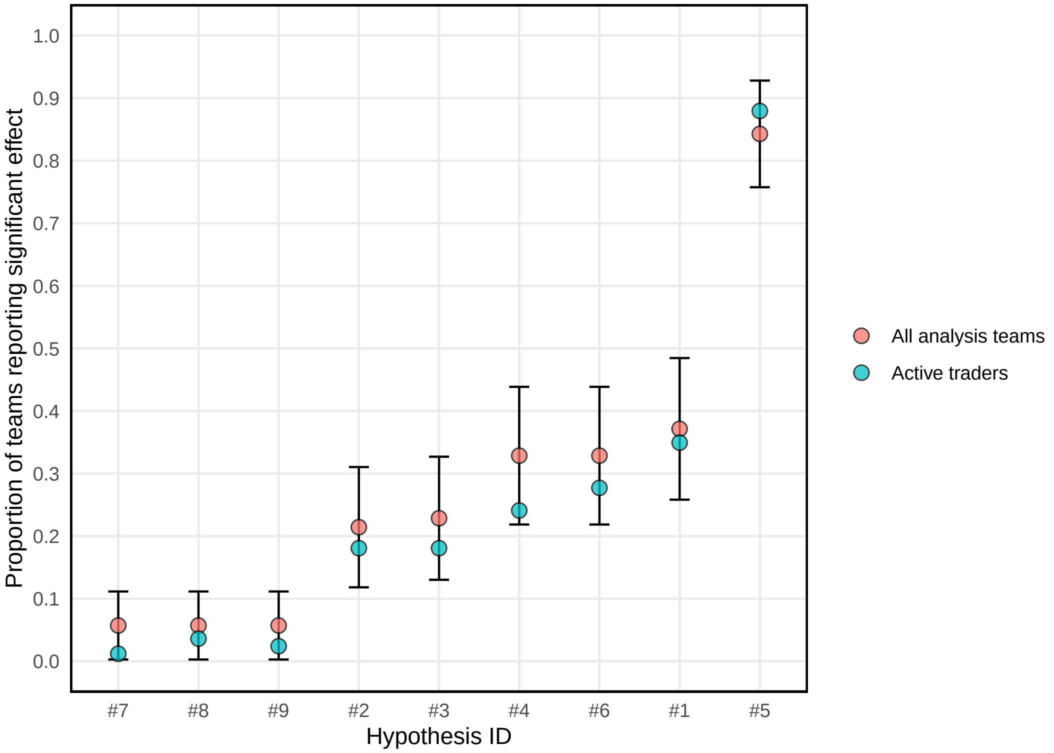


Supplementary Figure 11: The fraction of significant results is presented for each hypothesis, for all analysis teams (red) and separately for active traders only (green). The error bars represent the 95% confidence interval for the mean of all traders, computed using a normal approximation.

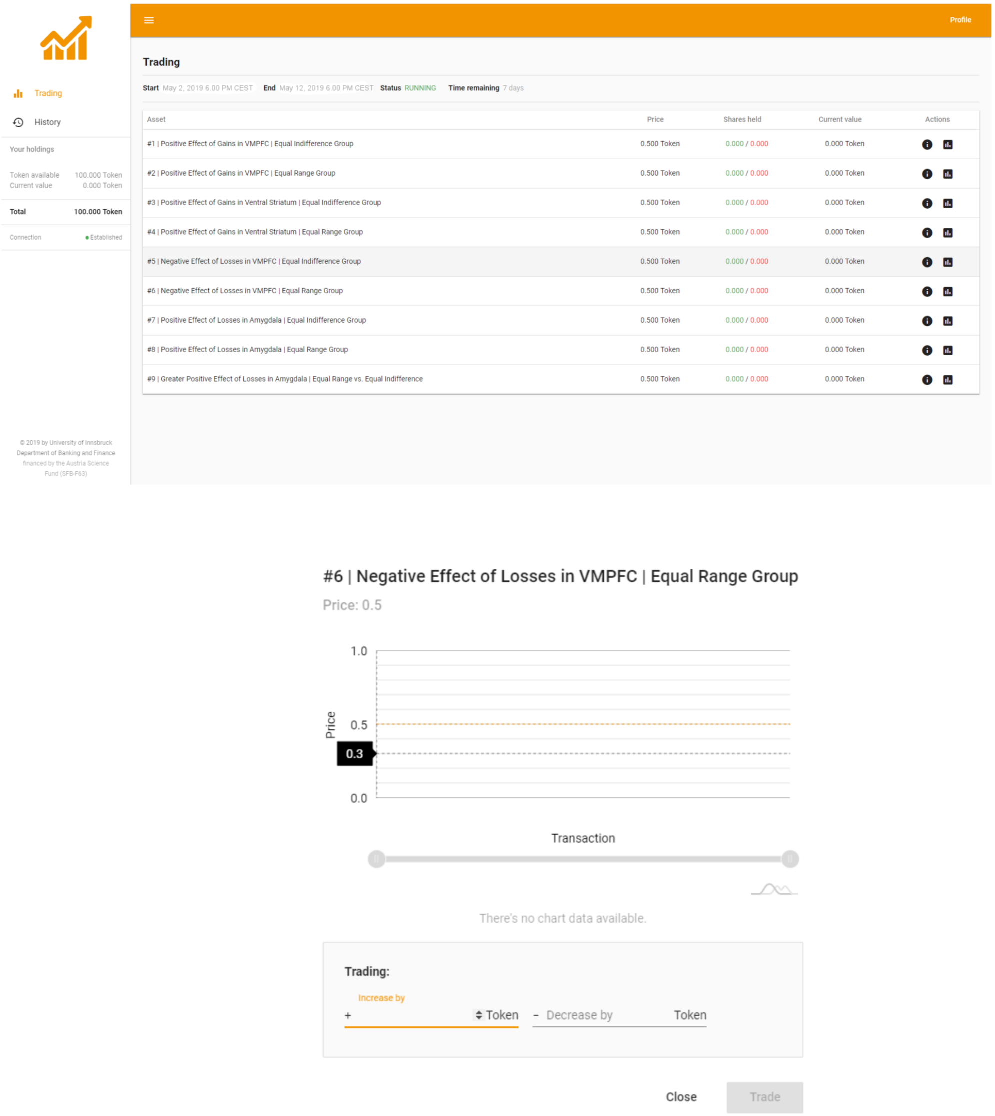


Supplementary Figure 12: Screenshots of the market overview (top panel) and the trading interface for a particular hypothesis (bottom panel). On the left hand side of the web-based interface, traders were informed about their current balance (i.e., the Tokens available) and the sum of the current value of the Tokens invested. The history tab showed a table of all transactions that have been submitted by the trader so far.

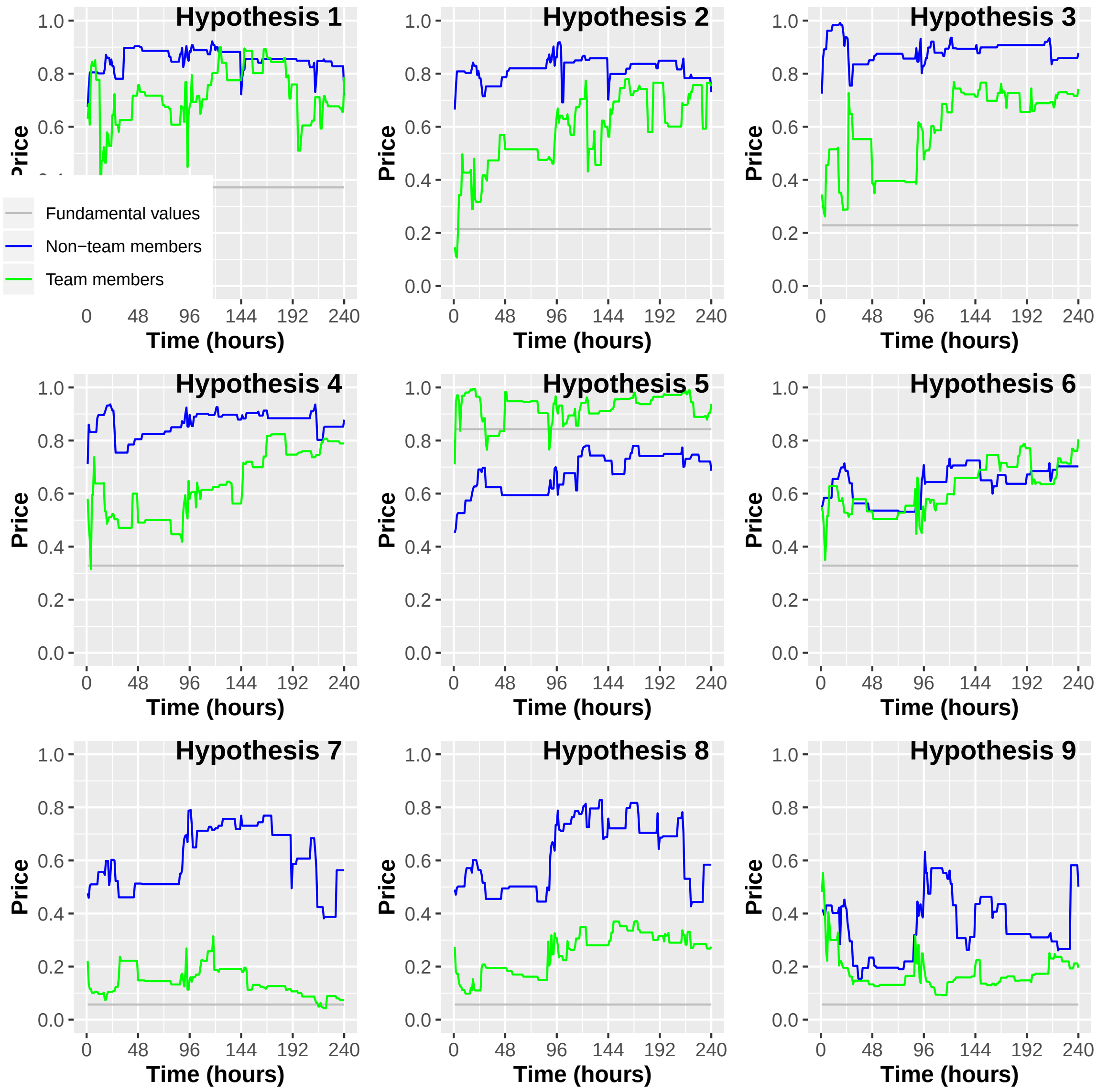


Supplementary Figure 13: Market prices for each of the nine hypotheses separated for the team members (green) and non-team members (blue) prediction markets. The figure shows the average prediction market prices per hour separated for the two prediction markets for the time the markets were open (10 days, i.e., 240 hours). The gray line indicates the actual share of analysis teams reporting a significant result for the particular hypothesis (i.e., the fundamental value).

| Team ID | 1 | 2 | 3 | 4 | 5 | 6 | 7 | 8 | 9 | fwhm | package | fmriprep | testing | movement_modeling |
|---|---|---|---|---|---|---|---|---|---|---|---|---|---|---|
| 4SZ2 | 7 | 5 | 6 | 6 | 9 | 9 | 7 | 8 | 7 | 6.65 | FSL | Yes | P | No |
| 1K0E | 7 | 9 | 6 | 6 | 8 | 7 | 7 | 6 | 9 | | Other | No | NP | Yes |
| R42Q | 5 | 5 | 6 | 6 | 6 | 6 | 7 | 8 | 8 | 12.73 | Other | No | P | Yes |
| Q58J | 9 | 9 | 9 | 9 | 9 | 9 | 9 | 9 | 9 | 16.24 | FSL | No | P | No |
| L1A8 | 8 | 5 | 7 | 7 | 8 | 8 | 3 | 8 | 3 | | SPM | No | P | Yes |
| 50GV | 10 | 10 | 10 | 10 | 10 | 10 | 10 | 10 | 10 | 10.26 | FSL | Yes | P | No |
| T54A | 5 | 9 | 2 | 6 | 9 | 9 | 5 | 5 | 5 | 12.28 | FSL | Yes | NP | No |
| R9K3 | 5 | 3 | 2 | 5 | 8 | 5 | 3 | 4 | 5 | 11.77 | SPM | Yes | P | Yes |
| P5F3 | 3 | 5 | 7 | 7 | 4 | 4 | 6 | 6 | 7 | 12.94 | FSL | No | P | Yes |
| IZ20 | 7 | 7 | 7 | 7 | 7 | 7 | 7 | 6 | 6 | 21.28 | Other | No | P | No |
| 9Q6R | 10 | 10 | 10 | 10 | 10 | 10 | 8 | 8 | 8 | 10.28 | FSL | No | P | Yes |
| XU70 | 4 | 5 | 8 | 9 | 9 | 9 | 6 | 8 | 8 | 7.17 | FSL | No | P | Yes |
| 80GC | 9 | 9 | 8 | 4 | 3 | 9 | 6 | 5 | 4 | 4.02 | AFNI | Yes | P | Yes |
| C22U | 8 | 7 | 5 | 8 | 9 | 8 | 8 | 8 | 8 | 11.16 | FSL | No | P | No |
| E6R3 | 5 | 5 | 7 | 3 | 4 | 4 | 7 | 7 | 7 | 9.28 | Other | Yes | Other | Yes |
| L9G5 | 5 | 4 | 4 | 6 | 10 | 10 | 9 | 9 | 7 | 7.22 | FSL | No | P | No |
| 2T6S | 8 | 9 | 6 | 6 | 10 | 9 | 7 | 8 | 10 | 14.93 | SPM | Yes | P | Yes |
| R5K7 | 6 | 8 | 8 | 7 | 9 | 7 | 8 | 8 | 7 | 12.06 | SPM | No | P | Yes |
| 98BT | 9 | 7 | 7 | 8 | 9 | 7 | 8 | 8 | 8 | 11.48 | SPM | No | P | Yes |
| L7J7 | 10 | 9 | 9 | 5 | 8 | 8 | 8 | 9 | 8 | 11.76 | SPM | Yes | P | Yes |
| O21U | 8 | 8 | 8 | 8 | 8 | 8 | 8 | 8 | 8 | 8.26 | FSL | Yes | P | Yes |
| UI76 | 10 | 6 | 10 | 10 | 10 | 6 | 10 | 10 | 5 | 6.60 | AFNI | Yes | P | Yes |
| E3B6 | 3 | 7 | 6 | 6 | 8 | 8 | 7 | 7 | 7 | 12.80 | SPM | Yes | P | Yes |
| DC61 | 5 | 1 | 5 | 2 | 9 | 5 | 5 | 5 | 5 | 9.58 | SPM | Yes | P | Yes |
| VG39 | 6 | 7 | 8 | 8 | 10 | 7 | 9 | 6 | 5 | | SPM | Yes | P | No |
| B5I6 | 10 | 10 | 5 | 5 | 10 | 6 | 8 | 7 | 6 | 9.84 | FSL | Yes | NP | Yes |
| X1Z4 | 8 | 6 | 4 | 4 | 9 | 5 | 4 | 4 | 4 | | Other | No | NP | Yes |
| 9U7M | 7 | 9 | 9 | 9 | 9 | 7 | 9 | 7 | 7 | 14.78 | Other | No | P | Yes |
| 08MQ | 8 | 6 | 8 | 6 | 7 | 7 | 7 | 7 | 6 | 13.14 | FSL | No | NP | Yes |
| 3C6G | 6 | 7 | 7 | 5 | 8 | 8 | 8 | 8 | 8 | 14.26 | SPM | No | P | Yes |
| 46CD | 9 | 8 | 5 | 8 | 9 | 8 | 9 | 9 | 5 | 10.92 | Other | No | P | Yes |
| 1P0Y | 8 | 8 | 1 | 1 | 8 | 8 | 5 | 5 | 5 | 9.13 | SPM | No | P | No |
| 43FJ | 3 | 3 | 5 | 5 | 10 | 10 | 10 | 10 | 10 | 10.66 | FSL | No | P | Yes |
| R7D1 | 4 | 7 | 5 | 5 | 9 | 5 | 8 | 9 | 8 | 8.93 | Other | Yes | NP | Yes |
| 16IN | 8 | 7 | 6 | 6 | 8 | 7 | 8 | 6 | 6 | | Other | Yes | Other | No |
| 6FH5 | 9 | 2 | 8 | 8 | 10 | 8 | 8 | 9 | 9 | 12.22 | SPM | No | P | Yes |
| 0H5E | 4 | 7 | 7 | 6 | 8 | 5 | 8 | 7 | 1 | 14.17 | SPM | No | P | No |
| V55J | 4 | 5 | 7 | 7 | 4 | 7 | 5 | 7 | 7 | 12.85 | SPM | No | P | No |
| 51PW | 8 | 8 | 8 | 8 | 8 | 8 | 6 | 6 | 7 | 11.15 | FSL | Yes | P | Yes |
| 4TQ6 | 7 | 9 | 10 | 9 | 7 | 8 | 10 | 10 | 9 | 14.88 | FSL | Yes | NP | No |
| I07H | 3 | 3 | 3 | 3 | 9 | 9 | 9 | 9 | 9 | 5.59 | Other | Yes | NP | No |
| 3PQ2 | 9 | 8 | 7 | 7 | 7 | 8 | 8 | 8 | 7 | 5.79 | FSL | No | P | Yes |
| L3V8 | 9 | 9 | 9 | 9 | 9 | 9 | 9 | 9 | 9 | 14.74 | SPM | No | P | No |
| K9P0 | 10 | 10 | 10 | 5 | 10 | 8 | 9 | 9 | 10 | 8.05 | AFNI | Yes | P | Yes |
| SM54 | 5 | 9 | 5 | 8 | 8 | 6 | 8 | 8 | 8 | 7.05 | Other | Yes | P | Yes |
| O03M | 3 | 8 | 8 | 2 | 8 | 7 | 7 | 7 | 7 | 3.47 | AFNI | Yes | NP | Yes |
| 0JO0 | 7 | 5 | 5 | 5 | 5 | 5 | 5 | 5 | 5 | 8.12 | Other | Yes | P | Yes |
| Q6O0 | 7 | 8 | 8 | 9 | 9 | 8 | 8 | 6 | 7 | 14.58 | SPM | Yes | P | Yes |
| 0I4U | 4 | 7 | 6 | 8 | 9 | 9 | 9 | 9 | 9 | 8.69 | SPM | No | P | Yes |
| X19V | 6 | 7 | 8 | 5 | 9 | 6 | 9 | 9 | 9 | 8.48 | FSL | Yes | P | Yes |
| X1Y5 | 6 | 6 | 7 | 7 | 8 | 6 | 8 | 8 | 8 | 8.69 | Other | Yes | NP | Yes |
| 0ED6 | 7 | 9 | 8 | 7 | 8 | 8 | 9 | 9 | 6 | 7.86 | SPM | No | P | Yes |
| U26C | 8 | 8 | 8 | 8 | 10 | 8 | 8 | 8 | 9 | 10.38 | SPM | Yes | P | Yes |
| C88N | 7 | 8 | 7 | 4 | 9 | 7 | 8 | 8 | 6 | 11.62 | SPM | Yes | P | No |
| 27SS | 4 | 6 | 7 | 7 | 7 | 7 | 6 | 8 | 4 | 11.37 | AFNI | No | P | Yes |
| O6R6 | 8 | 8 | 8 | 8 | 8 | 8 | 8 | 8 | 8 | 3.06 | FSL | Yes | NP | No |
| 94GU | 8 | 8 | 8 | 8 | 8 | 8 | 8 | 8 | 8 | 11.19 | SPM | No | P | Yes |
| 3TR7 | 2 | 2 | 3 | 4 | 8 | 5 | 8 | 6 | 5 | 17.40 | SPM | Yes | P | Yes |
| J7F9 | 9 | 8 | 9 | 7 | 9 | 7 | 9 | 9 | 9 | 14.88 | SPM | Yes | P | Yes |
| I52Y | 8 | 8 | 8 | 8 | 8 | 8 | 8 | 8 | 8 | 11.42 | FSL | No | NP | Yes |
| 0C7Q | 7 | 7 | 8 | 8 | 8 | 7 | 10 | 10 | 9 | 8.68 | Other | Yes | NP | Yes |
| 6VV2 | 8 | 8 | 8 | 6 | 9 | 7 | 8 | 7 | 6 | 7.20 | AFNI | No | P | Yes |
| B23O | 6 | 6 | 7 | 7 | 8 | 7 | 6 | 6 | 8 | 3.32 | FSL | Yes | NP | No |
| AO86 | 7 | 7 | 7 | 7 | 7 | 7 | 7 | 7 | 7 | 7.49 | Other | Yes | NP | Yes |
| 2T7P | 8 | 8 | 8 | 8 | 8 | 8 | 8 | 8 | 8 | 7.66 | Other | No | Other | Yes |
| 1KB2 | 6 | 6 | 8 | 8 | 5 | 5 | 8 | 8 | 7 | 13.06 | FSL | No | P | Yes |
| I9D6 | 7 | 7 | 7 | 7 | 1 | 7 | 7 | 6 | 7 | 6.21 | AFNI | No | P | Yes |
| UK24 | 4 | 4 | 4 | 4 | 4 | 4 | 4 | 4 | 4 | 10.76 | SPM | No | P | No |
| 5G9K | 7 | 7 | 7 | 7 | 7 | 7 | 7 | 7 | 7 | | SPM | Yes | P | Yes |
| 9T8E | 5 | 5 | 5 | 5 | 5 | 5 | 5 | 5 | 4 | 9.85 | SPM | Yes | NP | Yes |

Hypothesis number

Supplementary Table 1: Results submitted by analysis teams*. For each team (represented by its team ID, left column), the left section of the table represents the reported binary decision (green = yes, red = no) and how confident they were in their result (from 1 [not at all] to 10 [extremely]). The right section displays the information included for each team in the statistical model for hypothesis decisions. FWHM: estimated smoothing (full width at half-maximum). Teams with a blank value for the FWHM variable (estimated smoothness) were excluded from further analysis. Testing: P = parametric, NP = nonparametric. * It should be noted that three teams changed their decisions after the end of the project. Team L3V8 changed their decision regarding Hypothesis #6 from yes to no. Team VG39 changed their decisions regarding Hypotheses #3, #4 and #5 from yes to no. Team U26C changed their decision regarding Hypothesis #5 from yes to no. Results along the paper and in this table reflect the final results as they were reported at the end of the project (i.e., before this change), as prediction markets were based on those results.

Supplementary Table 2: Links to public NeuroVault collections of all analysis teams.

| Team ID | Link | Team ID.1 | Link.1 |
|---|---|---|---|
| 08MQ | https://neurovault.org/collections/4953/ | C88N | https://neurovault.org/collections/4812/ |
| 0C7Q | https://neurovault.org/collections/5652/ | DC61 | https://neurovault.org/collections/4963/ |
| 0ED6 | https://neurovault.org/collections/4994/ | E3B6 | https://neurovault.org/collections/4782/ |
| 0H5E | https://neurovault.org/collections/4936/ | E6R3 | https://neurovault.org/collections/4959/ |
| 0I4U | https://neurovault.org/collections/4938/ | I07H | https://neurovault.org/collections/5001/ |
| 0JO0 | https://neurovault.org/collections/4807/ | I52Y | https://neurovault.org/collections/4933/ |
| 16IN | https://neurovault.org/collections/4927/ | I9D6 | https://neurovault.org/collections/4978/ |
| 1K0E | https://neurovault.org/collections/4974/ | IZ20 | https://neurovault.org/collections/4979/ |
| 1KB2 | https://neurovault.org/collections/4945/ | J7F9 | https://neurovault.org/collections/4949/ |
| 1P0Y | https://neurovault.org/collections/5649/ | K9P0 | https://neurovault.org/collections/4961/ |
| 27SS | https://neurovault.org/collections/4975/ | L1A8 | https://neurovault.org/collections/5680/ |
| 2T6S | https://neurovault.org/collections/4881/ | L3V8 | https://neurovault.org/collections/4888/ |
| 2T7P | https://neurovault.org/collections/4917/ | L7J7 | https://neurovault.org/collections/4866/ |
| 3C6G | https://neurovault.org/collections/4772/ | L9G5 | https://neurovault.org/collections/5173/ |
| 3PQ2 | https://neurovault.org/collections/4904/ | O03M | https://neurovault.org/collections/4972/ |
| 3TR7 | https://neurovault.org/collections/4966/ | O21U | https://neurovault.org/collections/4779/ |
| 43FJ | https://neurovault.org/collections/4824/ | O6R6 | https://neurovault.org/collections/4907/ |
| 46CD | https://neurovault.org/collections/5637/ | P5F3 | https://neurovault.org/collections/4967/ |
| 4SZ2 | https://neurovault.org/collections/5665/ | Q58J | https://neurovault.org/collections/5164/ |
| 4TQ6 | https://neurovault.org/collections/4869/ | Q6O0 | https://neurovault.org/collections/4968/ |
| 50GV | https://neurovault.org/collections/4735/ | R42Q | https://neurovault.org/collections/5619/ |
| 51PW | https://neurovault.org/collections/5167/ | R5K7 | https://neurovault.org/collections/4950/ |
| 5G9K | https://neurovault.org/collections/4920/ | R7D1 | https://neurovault.org/collections/4954/ |
| 6FH5 | https://neurovault.org/collections/5663/ | R9K3 | https://neurovault.org/collections/4802/ |
| 6VV2 | https://neurovault.org/collections/4883/ | SM54 | https://neurovault.org/collections/5675/ |
| 80GC | https://neurovault.org/collections/4891/ | T54A | https://neurovault.org/collections/4876/ |
| 94GU | https://neurovault.org/collections/5626/ | U26C | https://neurovault.org/collections/4820/ |
| 98BT | https://neurovault.org/collections/4988/ | UI76 | https://neurovault.org/collections/4821/ |
| 9Q6R | https://neurovault.org/collections/4765/ | UK24 | https://neurovault.org/collections/4908/ |
| 9T8E | https://neurovault.org/collections/4870/ | V55J | https://neurovault.org/collections/4919/ |
| 9U7M | https://neurovault.org/collections/4965/ | VG39 | https://neurovault.org/collections/5496/ |
| AO86 | https://neurovault.org/collections/4932/ | X19V | https://neurovault.org/collections/4947/ |
| B23O | https://neurovault.org/collections/4984/ | X1Y5 | https://neurovault.org/collections/4898/ |
| B5I6 | https://neurovault.org/collections/4941/ | X1Z4 | https://neurovault.org/collections/4951/ |
| C22U | https://neurovault.org/collections/5653/ | XU70 | https://neurovault.org/collections/4990/ |

Supplementary Table 3: Description of teams excluded from the analyses of statistical maps.

| Team ID | Exclusion reason | Unthresholded maps excluded | Thresholded maps excluded |
|---|---|---|---|
| 1K0E | Used surface-based analysis (only provided data for cortical ribbon) | X | X |
| L1A8 | Not in MNI standard space | X | X |
| VG39 | Performed small volume corrected instead of whole-brain analysis | X | X |
| X1Z4 | Used surface-based analysis (only provided data for cortical ribbon) | X | X |
| 16IN | Values in the unthresholded images are not z / t stats | X | |
| 5G9K | Values in the unthresholded images are not z / t stats | X | |
| 2T7P | Used a method which does not create thresholded images (and are therefore not included in the analyses of the thresholded images) | | X |

Supplementary Table 4: Summary of mixed-effects logistic regression modeling of decision outcomes as a function of different factors including the hypothesis (1-9) and various aspects of statistical modeling including estimated spatial smoothing, use of fMRIprep preprocessed data, software package, multiple testing correction method, and use of movement modeling. For modeling details, see https://github.com/poldrack/narps/blob/master/ImageAnalyses/DecisionAnalysis.Rmd.

| Effects | Chi-squared | P value | Delta $R^2$ |
|---|---|---|---|
| Hypothesis | 185.390 | 0.000 | 0.350 |
| Estimated smoothness | 13.210 | 0.000 | 0.040 |
| Used fMRIPprep data | 2.270 | 0.132 | 0.010 |
| Software package | 13.450 | 0.004 | 0.040 |
| Multiple correction method | 7.500 | 0.024 | 0.020 |
| Movement modeling | 1.160 | 0.281 | 0.000 |

Supplementary Table 5: Variability in the number of activated voxels reported across teams.

| Hyp # | Minimum sig voxels | Maximum sig voxels | Median sig voxels | N empty images |
|---|---|---|---|---|
| 1 | 0 | 118181 | 1940 | 8 |
| 2 | 0 | 135583 | 8120 | 2 |
| 3 | 0 | 118181 | 1940 | 8 |
| 4 | 0 | 135583 | 8120 | 3 |
| 5 | 0 | 76569 | 6527 | 11 |
| 6 | 0 | 72732 | 167 | 25 |
| 7 | 0 | 147087 | 9383 | 8 |
| 8 | 0 | 129979 | 475 | 16 |
| 9 | 0 | 49062 | 266 | 29 |

Supplementary Table 6: Mean Spearman correlation between the unthresholded statistical maps for all pairs of teams and separately for pairs of teams within each cluster, for each hypothesis.

| Hyp | Correlation (mean) | Correlation (cluster1) | Cluster size (cluster1) | Correlation (cluster2) | Cluster size (cluster2) | Correlation (cluster3) | Cluster size (cluster3) |
|---|---|---|---|---|---|---|---|
| 1/3 | 0.394 | 0.670 | 50.000 | 0.680 | 7.000 | 0.095 | 7.000 |
| 2/4 | 0.521 | 0.736 | 43.000 | 0.253 | 14.000 | 0.659 | 7.000 |
| 5 | 0.485 | 0.777 | 41.000 | 0.329 | 20.000 | 0.342 | 3.000 |
| 6 | 0.259 | 0.442 | 47.000 | 0.442 | 12.000 | 0.156 | 5.000 |
| 7 | 0.487 | 0.851 | 31.000 | 0.466 | 25.000 | 0.049 | 8.000 |
| 8 | 0.302 | 0.593 | 36.000 | 0.256 | 23.000 | -0.044 | 5.000 |
| 9 | 0.205 | 0.561 | 47.000 | 0.568 | 8.000 | 0.106 | 9.000 |

Supplementary Table 7: Results from re-thresholding of unthresholded maps using uncorrected ($p < 0.001$, cluster size $k > 10$) and false discovery rate correction ($pFDR < 5\%$) and common anatomical regions of interest for each hypothesis. A team is recorded as having an activation (act.) if one or more significant voxels are found in the ROI. Results for coordinate-based meta-analysis (CBMA) and image-based meta-analysis (IBMA) for each hypothesis are also presented, each thresholded at $pFDR < 5\%$ as well.

| Hypothesis | N voxels in ROI | proportion of teams reporting act. | proportion of teams w/ act. ($p < 0.001$, $k > 10$) | proportion of teams w/ act. (FDR) | CBMA (n voxels in ROI) | IBMA (n voxels in ROI) |
|---|---|---|---|---|---|---|
| 1 | 3402.000 | 0.371 | 0.734 | 0.594 | 1184.000 | 0.000 |
| 2 | 3402.000 | 0.214 | 0.391 | 0.766 | 144.000 | 7.000 |
| 3 | 173.000 | 0.229 | 0.156 | 0.344 | 56.000 | 0.000 |
| 4 | 173.000 | 0.329 | 0.234 | 0.609 | 65.000 | 7.000 |
| 5 | 3402.000 | 0.843 | 0.906 | 0.859 | 2815.000 | 2101.000 |
| 6 | 3402.000 | 0.329 | 0.562 | 0.359 | 2265.000 | 39.000 |
| 7 | 672.000 | 0.057 | 0.062 | 0.172 | 0.000 | 0.000 |
| 8 | 672.000 | 0.057 | 0.016 | 0.125 | 4.000 | 0.000 |
| 9 | 672.000 | 0.057 | 0.047 | 0.094 | 2.000 | 0.000 |

Supplementary Table 8: Prediction market results. The table summarizes the prediction market results for each of the nine ex-ante hypotheses, separated for the team members and non-team members prediction markets. FV indicates the fundamental value, i.e., the actual fraction of teams reporting significant results for the particular hypothesis. 95% CI refers to the 95% confidence interval corresponding to the fundamental value. Market belief refers to the final prediction market price (i.e. markets predictions) and within CI indicates whether the market beliefs are within or outside the 95% confidence interval. Note that this is not a formal hypothesis test as it does not take into account the uncertainty in the final prediction market prices, given that we have no measure of the standard error for the aggregated market prediction for each specific hypothesis. Thus, only for the single prediction that is within the 95% confidence interval (Hypothesis #7) it is clear that the aggregated belief does not differ significantly from the fundamental value.

| Hyp # | FV | CI | Non-teams FV | Teams FV |
|---|---|---|---|---|
| 1 | 0.370 | [0.26-0.48] | 0.727 * | 0.814 * |
| 2 | 0.210 | [0.12-0.31] | 0.73 * | 0.753 * |
| 3 | 0.230 | [0.13-0.33] | 0.881 * | 0.743 * |
| 4 | 0.330 | [0.22-0.44] | 0.882 * | 0.789 * |
| 5 | 0.840 | [0.76-0.93] | 0.686 * | 0.952 * |
| 6 | 0.330 | [0.22-0.44] | 0.685 * | 0.805 * |
| 7 | 0.060 | [0.00-0.11] | 0.563 * | 0.073 |
| 8 | 0.060 | [0.00-0.11] | 0.584 * | 0.274 * |
| 9 | 0.060 | [0.00-0.11] | 0.476 * | 0.188 * |

Supplementary Table 9: Consistency of traders holdings and team results. The top section of the table reports Spearman rank correlations (s) between traders final holdings and the binary result reported by their team and the corresponding p-value for each of the nine hypotheses. The lower section reports the share of traders holdings that are consistent with the results reported by their team. In particular, consistent refers to positive (negative) holdings if the team reported a significant (non-significant) result. z- and p-values refer to Wilcoxon signed-rank tests for the share of consistent holdings being equal to 0.5. Avg. holdings if (in)consistent refer to the mean final holdings, separated for consistent and inconsistent traders.

| Hypothesis # | 1 | 2 | 3 | 4 | 5 | 6 | 7 | 8 | 9 |
|---|---|---|---|---|---|---|---|---|---|
| Spearman rho | 0.580 | 0.560 | 0.580 | 0.640 | 0.470 | 0.740 | 0.230 | 0.370 | 0.310 |
| p-value | 0.000 | 0.000 | 0.000 | 0.000 | 0.000 | 0.000 | 0.104 | 0.007 | 0.020 |
| Share of consistent holdings | 0.710 | 0.680 | 0.700 | 0.800 | 0.890 | 0.740 | 0.800 | 0.800 | 0.750 |
| Z (signed rank test) | 3.400 | 2.780 | 2.820 | 4.240 | 6.810 | 3.240 | 4.340 | 4.340 | 3.640 |
| p-value (signed rank test) | 0.000 | 0.003 | 0.002 | 0.000 | 0.000 | 0.001 | 0.000 | 0.000 | 0.000 |
| Average holdings if consistent | 5.610 | 21.140 | 25.800 | 13.110 | -115.500 | 7.310 | 34.610 | 24.230 | 23.540 |
| Average holdings if inconsistent | 1.040 | -6.900 | -8.030 | 0.030 | 18.260 | 1.580 | -14.630 | -8.290 | -11.610 |

Supplementary Table 10: Links to shared analysis codes of some of the analysis teams.

| Team ID | Link to shared analysis codes |
|---|---|
| 16IN | https://github.com/jennyrieck/NARPS |
| 2T7P | https://osf.io/3b57r |
| E3B6 | DOI: 10.5281/zenodo.3518407 |
| Q58J | https://github.com/amrka/NARPS_Q58J |

Supplementary Table 11: Market details. The table depicts additional data for each of the nine hypotheses, separated for the team members and non-team members prediction markets. Tokens invested indicates the average number of token invested per transaction and Volume (Shares) refers to the mean number of shares bought or sold per transaction. Transactions describes the overall number of transactions recorded and No. of Traders refers to the number of traders who bought or sold shares of the particular asset at least once.

| Hyp # | Tokens invested (Non-teams) | Volume (Non-teams) | # Traders (Non-teams) | # Transactions (Non-teams) | Tokens invested (Teams) | Volume (Teams) | # Transactions (Teams) | # Traders (Teams) |
|---|---|---|---|---|---|---|---|---|
| 1 | 8.568 | 20.175 | 55 | 139 | 12.643 | 25.671 | 213 | 64 |
| 2 | 10.510 | 22.544 | 53 | 98 | 11.632 | 22.908 | 171 | 58 |
| 3 | 12.818 | 24.709 | 58 | 132 | 7.773 | 15.837 | 141 | 52 |
| 4 | 11.134 | 20.397 | 49 | 112 | 8.126 | 15.479 | 127 | 52 |
| 5 | 6.873 | 14.636 | 38 | 71 | 14.480 | 30.760 | 244 | 76 |
| 6 | 6.806 | 12.663 | 35 | 72 | 8.097 | 16.676 | 134 | 46 |
| 7 | 7.990 | 15.209 | 41 | 98 | 7.131 | 15.864 | 160 | 52 |
| 8 | 8.791 | 19.072 | 45 | 91 | 7.085 | 14.598 | 141 | 52 |
| 9 | 10.427 | 21.118 | 50 | 131 | 9.506 | 18.812 | 178 | 56 |

Supplementary Table 12: Panel regressions. The table summarizes the results of pre-registered fixed-effects panel regressions of the predictions absolute errors (i.e., the absolute deviation of the market price from the fundamental value) on an hourly basis (average price of all transactions within an hour) on time and prediction market indicators. Standard errors are computed using a robust estimator.

| Effect | Beta (full model) | t (full model) | p (full model) | Beta (no interaction) | t (no interaction) | p (no interaction) |
|---|---|---|---|---|---|---|
| Intercept | 0.440 | 64.120 | 0.000 | 0.410 | 74.610 | 0.000 |
| Time | 0.000 | 3.380 | 0.001 | 0.000 | 12.480 | 0.000 |
| Teams | -0.290 | -29.500 | 0.000 | -0.220 | -45.350 | 0.000 |
| Time X Teams | 0.000 | 7.780 | 0.000 | | | |
| ——— | | | | | | |
| Adjusted R-squared | | | 0.350 | | | 0.340 |
